# Mutations to the HCoV-229E spike have counterbalancing effects on serum antibody neutralization and receptor binding

**DOI:** 10.64898/2026.02.22.707297

**Authors:** Sheri Harari, Rachel T. Eguia, Bernadeta Dadonaite, Caelan E. Radford, Cameron Stewart, David Veesler, Jesse D. Bloom

## Abstract

Human coronavirus 229E (HCoV-229E) is an endemic pathogen that causes repeated “common-cold” infections throughout life. Like other coronaviruses, it accumulates spike mutations that erode antibody immunity and enable reinfection. Here, we use pseudovirus deep mutational scanning to measure how mutations to the HCoV-229E spike affect its cell entry function, binding to its human APN receptor, and neutralization by human sera with a range of sensitivities to erosion by viral evolution. We find that both receptor binding and serum neutralization are affected by mutations across spike, including many that modulate these properties by affecting the balance of up versus down conformations of the spike receptor-binding domain (RBD). In particular, some mutations increase both receptor binding and serum neutralization by shifting the RBD to a more up conformation, suggesting that the HCoV-229E spike has evolved to shield key RBD neutralizing epitopes at the cost of less efficient receptor binding.

## Introduction

The COVID-19 pandemic was caused by the introduction into humans of a novel coronavirus to which there was no pre-existing immunity. As population immunity accumulated, new SARS-CoV-2 variants began to evolve with spike mutations that enabled them to erode pre-existing antibody immunity to re-infect people, albeit usually with reduced disease severity^1–4^. This transition underscores the importance of understanding how coronaviruses evolve after they become endemic in humans, where they are under selective pressure to acquire spike mutations promoting immune evasion without disrupting the molecular functions required for infection and transmission^5,6^.

A natural comparison is provided by the four endemic seasonal human coronaviruses (HCoV-229E, HCoV-OC43, HCoV-NL63, and HCoV-HKU1), which have long circulated in humans and typically cause mild-to-moderate upper respiratory illness^7–11^. A typical person is reinfected with each of these endemic coronaviruses multiple times due to a combination of waning immunity and evolution of the viral spike to erode neutralization by pre-existing antibodies^5,6,12–14^. However, it remains unclear how the long-term evolution of the viral spike balances the acquisition of mutations that affect antigenicity with the need to maintain essential spike properties such as receptor binding and the ability to mediate cell entry^15–18^.

Here we examine these questions by studying the spike of human coronavirus 229E (HCoV-229E), which was the first human coronavirus to be identified (in 1966) and likely has been endemic in humans from long before its identification^12,14,19,20^. Human individuals are typically infected with HCoV-229E every ∼3-5 years, and its spike protein is known to undergo antigenic evolution to erode recognition by antibodies^5,6,12,13^.

The HCoV-229E spike binds to human aminopeptidase N (hAPN) as its receptor^21,22^ and then mediates viral entry by fusing the viral and cell membranes^16,17,23^; spike is also the primary target of neutralizing antibodies^24–26^. Spike must therefore balance strong functional constraints, including proper folding, protease activation, conformational rearrangements, and fusion, with intense immune selection that drives antigenic drift^5,6,27,28^.

Here we elucidate at the molecular level how HCoV-229E evolution has balanced these constraints and pressures on its spike during evolution. We do this by using pseudovirus deep mutational scanning platform to measure the effects of nearly all mutations to the spike of an historical human strain from 1984 on the key properties of cell entry function, receptor binding, and neutralization by human sera. Our work defines the constraints that have shaped HCoV-229E spike evolution, and uncover how evolution counterbalances the effects on receptor binding and antibody neutralization of mutations that modulate RBD up/down conformation.

## Results

### Deep mutational scanning measurement of how all mutations to HCoV-229E spike affect its cell entry function

To understand constraints that govern the evolution of the HCoV-229E spike, we experimentally measured the effects of all individual amino-acid mutations to this protein on some of its key phenotypes. We chose the spike of a strain from 1984 (Fig. S1A) to enable comparison of our measurements on this historical strain to subsequent virus evolution.

To measure the effects of mutations, we used pseudovirus deep mutational scanning^29^. In brief, this approach involves generating libraries of pseudotyped lentiviral particles that each encode a single spike protein variant that is also displayed on the virion surface. Each variant is linked to a unique nucleotide barcode that identifies the spike sequence encoded by that particle (Fig. S1B). To make high-throughput measurements by deep mutational scanning, these barcoded spike-pseudotyped lentiviral particles are generated in a pool and used to perform a single massively parallel experiment that quantifies the infectivity of each spike variant by sequencing of its barcode (Fig. S1C). The pseudotyped lentiviral particles (pseudoviruses) can only undergo a single round of cell entry since they do not encode any of the essential viral genes other than spike, and therefore provide a safe way to measure the effects of mutations of viral proteins at biosafety-level-2. For these experiments, we mutagenized the full ectodomain and most of the transmembrane domain of spike (sites 18-1140; throughout this manuscript we use HCoV-229E RefSeq NC_002645 numbering). To improve pseudovirus titers, we deleted the last 19 residues of the cytoplasmic tail^5,30–32^. We created two independent barcoded pseudovirus libraries, which contained 90,644 and 96,532 barcoded variants and covered 96% and 97% of all possible amino-acid mutations, respectively (Fig. S1D). Most barcoded spike variants had a single amino-acid mutation, but some had no mutations or multiple mutations (Fig. S1E).

We first measured the ability of each spike variant to mediate virion entry into 293T cells that we had engineered to constitutively express the HCoV-229E receptor human aminopeptidase N (hAPN)^21,22^ and the activating protease TMPRSS2^33,34^ (Fig. S2A-B). The cell entry of each variant was determined by comparing the barcode counts for that variant in a condition where pseudovirus entry was dependent on spike versus a control condition where it was mediated by VSV-G (Fig. S2C), quantifying cell entry as the log2 of ratio of that variant’s barcode counts in the spike-mediated versus VSV-G-mediated entry experiments after normalizing to the barcode counts for unmutated spike. Cell entry scores of zero indicate mutations have no effect on cell entry, whereas negative cell entry scores indicate impaired cell entry. As expected, spike variants with only synonymous mutations had cell entry scores of close to zero, variants with stop-codon mutations had highly negative scores, and variants with amino-acid mutations had a range of cell entry scores (Fig. S2D). To determine the effects of individual amino-acid mutations on cell entry, we deconvolved the scores for both single- and multi-mutant variants using a global epistasis model^35,36^. Measurements of mutation effects on cell entry were highly correlated both between technical replicate measurements made using the same library and across the two independent libraries (Fig. S2E).

The full set of cell entry measurements is shown in Fig. S3 and in interactive form at https://dms-vep.org/229E_spike_1984_DMS/cell_entry.html. Overall, the functional constraint on mutations varies across the spike (Fig. 1A,B and S3). On average, mutations are more deleterious in the S_2_ subunit of spike compared to the S_1_ subunit, although there are regions of mutational constraint and tolerance in both subunits (Fig. 1A-E). Within S_1_, the NTD is generally more tolerant of mutations than the RBD (Fig. 1A,B). There is also substantial functional constraint in the C-terminal region of S_1_ (approximately sites 446–564), which has been proposed to stabilize the prefusion spike trimer^37^. In S_2_, there is especially strong constraint for the interior-facing residues within the three core helices (HR1, CH, and UH), the S_2_’ cleavage site region (SCR; sites 686–711), the fusion peptide region (FP; sites 741–781^38^), and the HR1–CH junction loop (sites 865–871) (Fig. 1A-F). The complete intolerance for mutations at the S_2_’ cleavage site concurs with the key role of this residue for fusion activation^39–45^. We hypothesize that the constraint on the HR1–CH loop could be in part due to its proposed role in modulating RBD down versus up positioning^37,46^, consistent with its proximity to the RBD in the prefusion trimer, or due to the complete refolding of this region to form a continuous helix in postfusion spike^38^ (Fig. 1D).

**Figure 1.**
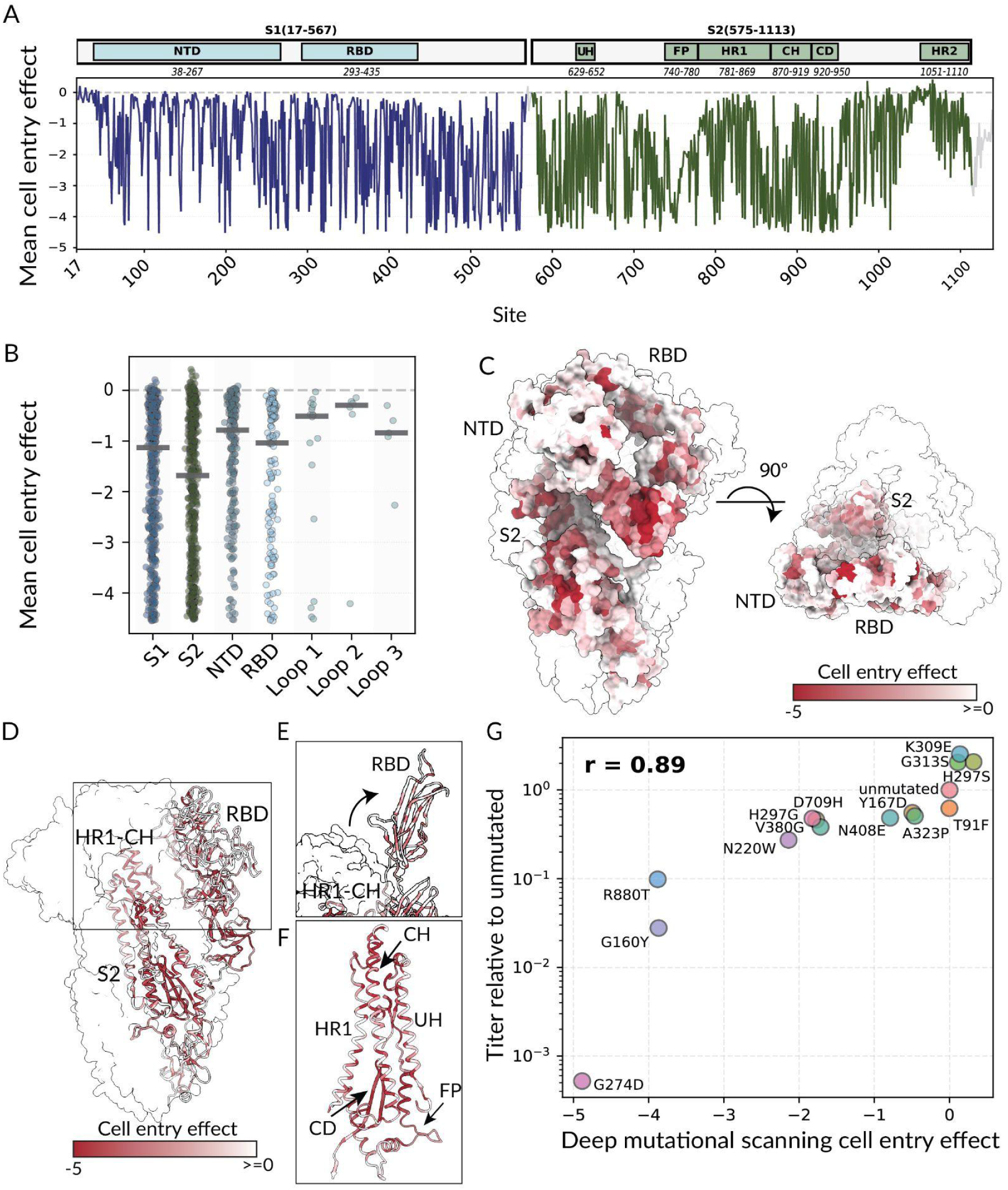
Effects of HCoV-229E spike mutations on cell entry. (A) Mean effect of mutations at each site in spike on cell entry as measured by deep mutational scanning (NTD, N-terminal domain; RBD, receptor binding domain; UH, upstream helix; FP, fusion peptide; HR1, heptad repeat 1; CH, central helix; CD, connector domain; HR2, heptad repeat 2). See https://dms-vep.org/229E_spike_1984_DMS/htmls/cell_entry_func_effects.html for an interactive plot showing the effects of mutations on cell entry in more detail. (B) Distribution of the mean effects of all mutations at each site in different regions of spike (S_1_, S_2_, NTD, RBD, and the three receptor binding loops). (C) The spike heterotrimer (PDB 7CYC^46^) with one monomer colored by the mean effect of all mutations at each site on cell entry, and the other two monomers shown in white. Darker red indicates mutations impair cell entry. (D) A different view of spike with the monomer colored by cell entry effects shown in a cartoon. The HR1-CH loop and RBD are labeled to demonstrate their structural proximity. (E) Zoomed in view of the spike RBD in the up conformation (PDB 8WDE^38^) again with one monomer colored by cell entry effects. (F) Zoomed in view of main S_2_ domains colored by cell entry effects (PDB 7CYC^46^). (G) Validation assays showing the correlation between the fold-change in titer of individually generated pseudotyped lentiviral particles carrying the indicated mutations versus the mutation effects on cell entry measured by deep mutational scanning. The titers are the mean of two biological replicates for each mutant, and r indicates the Pearson correlation.

To validate the deep mutational scanning measurements, we chose 14 mutations that spanned a range of effects on cell entry and generated individual spike-pseudotyped lentiviral particles with each mutation. The titers of the individual mutants were highly correlated with the mutation effects measured in the deep mutational scanning (Pearson R of 0.89; Fig. 1G).

### Comparison of measured effects of mutations on cell entry to natural HCoV-229E spike evolution

Among natural HCoV-229E strains, the most variable region of spike is the three loops in the RBD that contact the hAPN receptor (Fig. 2A and prior work^5,47^). The high variability of these loops is likely driven by immune pressure for the virus to evolve to erode antibody neutralization^5^, as the most potent neutralizing antibodies to coronaviruses tend to target the RBD and compete with its ability to bind receptor^25,48–51^. However, despite the high sequence variability in these binding loops, different HCoV-229E strains retain the ability to bind strongly to hAPN^52^.

**Figure 2.**
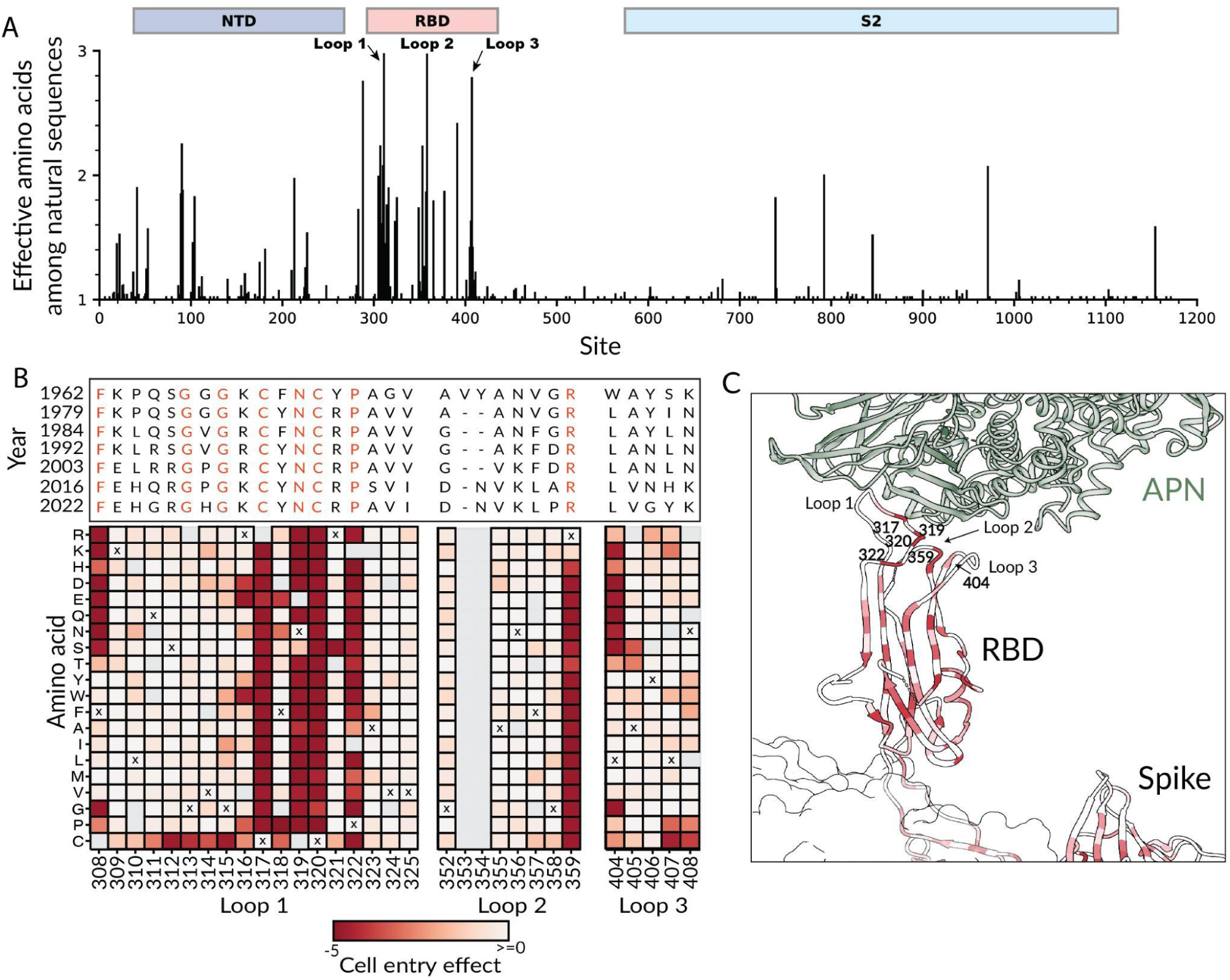
Natural variation of the RBD binding loops of HCoV-229E versus effects of mutations on cell entry measured in deep mutational scanning. (A) Per-site diversity across spike summarized as the effective number of amino acids among all publicly available NCBI HCoV-229E spike sequences. Domains are annotated above the plot, including the three RBD binding-loops. See https://dms-vep.org/229E_spike_1984_DMS/rbd_loops.html for an interactive version of this plot. (B) Effects of mutations to the RBD binding loops on cell entry as measured in the deep mutational scanning shown alongside an alignment of the loop sequences in HCoV-229E sequences from various years; note that the 1962 strain was extensively passaged in the lab prior to sequencing. Each square in the heatmap is colored according to the effect of that mutation on cell entry (red indicates impaired entry, white indicates no effect), and the “x” for each site indicates the amino-acid identity in the 1984 HCoV-229E spike used in the deep mutational scanning. Gray indicates no measurement was made for that mutation; note that there is an insertion of two residues after site 352 in the 1962 spike relative to the spike used in the deep mutational scanning. Conserved residues in the alignment are colored red. (C) Zoomed in view of the HCoV-229E spike RBD bound to hAPN (PDB 8WDE^47^) colored by the average effect of all mutations at each site on cell entry. The binding loops and key conserved sites are labeled.

Although many sites in the HCoV-229E RBD binding loops are highly variable among natural sequences, a subset of sites in these loops are conserved across HCoV-229E sequences from over the last six decades (Fig. 2B). Our deep mutational scanning provides an explanation for the conservation of these sites: at the conserved binding-loop sites most or all mutations are measured to strongly impair cell entry, whereas at the variable sites many mutations can be tolerated without strongly affecting cell entry (Fig. 2B). In particular, our deep mutational scanning shows that most mutations at Cys317, Asn319, Cys320, Pro322, and Arg359 strongly impair cell entry, and these sites are conserved among HCoV-229E from different timeframes (Fig. 2B). Structurally, these sites map to the center of the receptor binding interface (Fig. 2C), and have an important role in maintaining the receptor binding motif. For example, Cys317 and Cys320 form a disulfide bond that stabilizes the loop structure and Asn319 is a receptor-contact residue^11,18^. Leu404, another site that is highly conserved in natural sequences and strongly constrained in our data, forms a polar contact with hAPN^37^. Overall, these results suggest that extensive immune pressure during HCoV-229E evolution has led to sequence variation at nearly every site in the hAPN binding loops that can functionally tolerate mutations, but that some sites remain conserved due to high functional constraint.

### Mutations both in and outside the RBD affect binding to hAPN receptor

Mutations can affect spike-mediated cell entry by modulating several biochemical properties including protein folding, fusogenicity, and receptor binding. To define the constraints on the HCoV-229E spike in greater detail, we used our deep mutational scanning libraries to directly measure how mutations affect binding to the hAPN receptor. We did this by leveraging the previously established fact that inhibition of pseudovirus infection by soluble receptor is directly proportional to the binding affinity for the viral spike protein ^53–56^. Specifically, we incubated the pseudovirus libraries with a range of concentrations of soluble dimeric hAPN, also including VSV-G–pseudotyped particles with known barcodes as an internal standard to convert sequencing counts into the fraction of infectivity retained^29,53^ (VSV-G neutralization is not affected by hAPN) (Fig. S4A-B). This assay provides a quantitative readout of how each spike mutation affects hAPN binding: variants with stronger hAPN binding are neutralized more potently by soluble hAPN, whereas variants with reduced binding are less neutralized by soluble hAPN (Fig. S4A). Mutations can affect the overall spike-receptor binding measured in this assay by two mechanisms: altering the direct affinity of the RBD for hAPN, or modulating the up/down conformation of the RBD in the context of the full spike thereby affecting its accessibility for hAPN binding^53,54^. This assay can only measure the impact on hAPN binding of mutations that retain at least some minimal cell entry function, as the assay readout requires pseudovirus infection of target cells, therefore mutations that strongly impair cell entry are filtered out of the following analysis (Fig. S5).

Binding of the HCoV-229E spike to hAPN was affected both by mutations at sites in the RBD that contact hAPN and by some mutations elsewhere in spike, including RBD sites distal from the receptor, parts of the NTD, and some other regions of S_1_ and in S_2_ (Fig. 3A–D, Fig. S5, and interactive plots at https://dms-vep.org/229E_spike_1984_DMS/APN_binding.html). Mutations that disrupt known N-linked glycosylation motifs (N-X-S/T)^47^ are often strongly deleterious for cell entry; mutations that add glycosylation motifs but are tolerated for cell entry sometimes still affect hAPN binding (Fig. S6). As expected, many mutations at sites close to the receptor interface in the RBD, markedly reduced hAPN binding, consistent with disruption of direct receptor binding by the RBD^37,47^ (Fig. 3A, orange line, and Fig. 3B-D). However, binding is also affected by some mutations at sites far from the hAPN-RBD interface (Fig. 3A, gray and blue lines, Fig. 3B-D), especially at sites in parts of the RBD distal from APN and in the NTD. For example, mutations at NTD sites 87-93 generally increased hAPN binding, whereas mutations at NTD sites 147-152 mostly decreased binding (Fig. 3C, Fig. S5). Notably, these NTD sites are distant from the RBD receptor-binding loops and are not expected to contact hAPN when engaged to the spike based on available structural data, suggesting an indirect influence on receptor engagement (Fig 3B-C).

**Figure 3.**
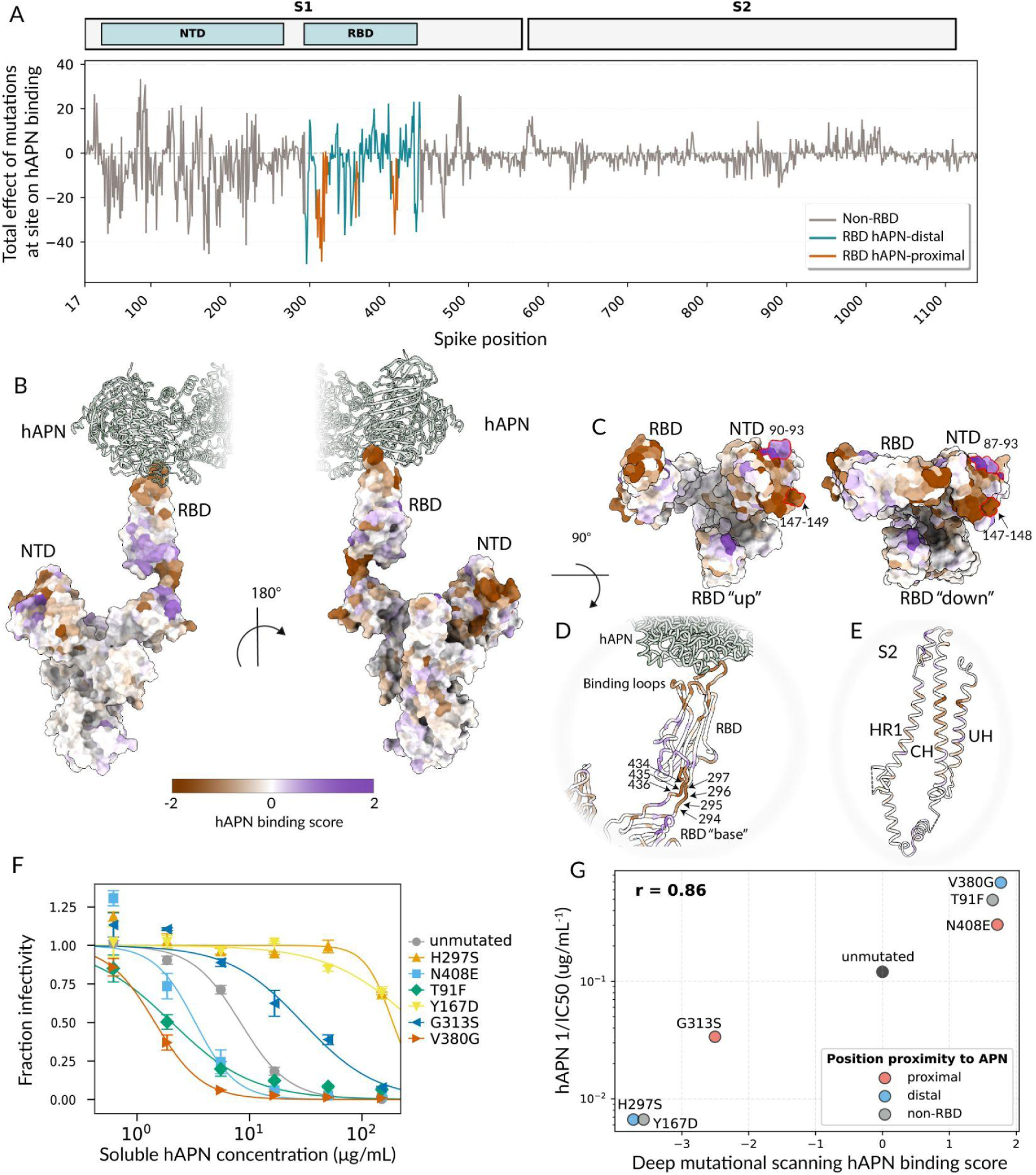
Mutations in both receptor contact sites and other regions of spike affect binding to hAPN as measured by pseudovirus neutralization by soluble hAPN. (A) Total effect of mutations at each site on hAPN binding by the HCoV-229E spike. Sites are colored based on proximity to hAPN, where proximal sites are defined as RBD sites within 15Å. A positive effect indicates increased receptor binding and a negative effect indicates decreased receptor binding. (B) One spike monomer (PDB 8WDE^47^) in complex with hAPN, colored by the total effect of all mutations at each site of receptor binding. Purple indicates increased binding and brown indicates decreased binding.(C) Top view of the spike structures with RBD “down” (PDB 6U7H^37^ and RBD “up” PDB 8WDE^47^) colored by the total effect of all mutations at each site on receptor binding. Sites with high receptor binding effects in the NTD are highlighted with a red outline as an example. (D-E) The spike structure in complex with hAPN shown as cartoon and colored as in B and C. (F) Validation assays showing the neutralization of lentiviral particles pseudotyped with HCoV-229E spike carrying the indicated mutations by soluble hAPN. (G) Correlation of the inverse IC_50_ from the validation assays in panel F versus the effect of that mutation on receptor binding measured in the deep mutational scanning. The plot shows the mean of two biological replicates for each mutant, and r indicates the Pearson correlation. See https://dms-vep.org/229E_spike_1984_DMS/APN_binding.html for interactive plots showing more details about how mutations affect hAPN binding

The mutations at RBD sites proximal to the hAPN binding interface probably affect receptor binding by directly modulating the RBD’s affinity for hAPN, but the mutations distal from the hAPN binding site likely affect the RBD’s up/down motion. Specifically, coronavirus spike RBDs can be positioned in either a down conformation, that masks the receptor-binding motif/loops, or an up conformation which exposes that motif. As a result, receptor binding requires the RBD to be in the up conformation^28,50,57,58^. Recent studies on spikes from other coronaviruses have shown that mutations outside the receptor-binding motif can have appreciable effects on overall spike receptor engagement by modulating the balance of up versus down RBD conformation^53,54,59–61^. Non-RBD mutations and hAPN-distal RBD mutations that affect receptor binding tend to be at sites that could affect the RBD’s up/down motion. For instance, mutations to the linkers directly N- and C-terminal to the RBD (sites 294-297 and 434-436, which are away from the receptor-binding loops; Fig. 3D, Fig. S5) strongly affect receptor binding, likely by modulating the dynamic of the RBD down to up movements^47^. A similar mechanism may apply to some S_2_ mutations: recent on-virion structural analysis shows that prefusion HCoV-229E spikes can exist in S_2_-compact and S_2_-loose conformations, with RBD up state being more enriched in the S_2_-loose conformation^62^. We detected some S_2_ mutations at sites in the three core helices (UH, HR1, CH) that directly impact transition between loose and compact state^46^ to affect hAPN binding (Fig. 3E, Fig. S5), likely by indirectly altering the balance of RBD up versus down conformations.

To independently validate the deep mutational scanning measurements of the effect of spike mutations on hAPN binding, we selected six substitutions spanning a range of effects, generated individual pseudoviruses expressing each mutant spike, and tested their neutralization by soluble hAPN. This set included mutations in the RBD proximal to the hAPN-binding interface (G313S, N408E), mutations in the RBD distal from the hAPN-binding interface (H297S, V380G), and two NTD mutations (T91F, Y167D), chosen so that within each region one mutation increased and one mutation decreased receptor binding in the deep mutational scanning (Fig. S4C and S5). The individual pseudovirus neutralization assays validated the deep mutational scanning measurements of the effects of these mutations on neutralization by soluble hAPN (Fig. 3G-F), confirming that mutations in multiple regions of spike can affect full-spike receptor binding as quantified by soluble hAPN neutralization.

To compare the relative effects of receptor-proximal RBD mutations to receptor-distal mutations on receptor binding for HCoV-229E to another coronavirus, we examined previously published deep mutational scanning measurements of how mutations to the SARS-CoV-2 KP.3.1.1 or XBB.1.5 spike affected receptor binding as quantified by pseudovirus neutralization by soluble monomeric human ACE2^53,54^. A caveat of this comparison is that our HCoV-229E deep mutational scanning encompassed nearly all amino-acid mutations, whereas the KP.3.1.1 deep mutational scanning encompassed all RBD mutations, but only naturally observed mutations or mutations otherwise thought to be tolerated at non-RBD sites, while the XBB.1.5 deep mutational scanning encompassed only naturally observed mutations or mutations otherwise thought to be tolerated at all sites. Given that S_2_ mutations were less well represented than S_1_ mutations in the SARS-CoV-2 spike deep mutational scanning data, we restricted this analysis solely to S_1_ mutations. Mutations at RBD sites that are distant from the receptor-binding loops tend to have a larger relative effect on receptor binding for HCoV-229E than observed for either SARS-CoV-2 variants (Fig. 4). Therefore, mutations that affect RBD up-down conformation are relatively more impactful on receptor binding for the HCoV-229E spike compared to the spikes of recent SARS-CoV-2 variants. This observation is consistent with various structural studies suggesting that the prefusion HCoV-229E spike trimer typically adopts a conformation with all RBDs in the closed state^37,47,62,63^, whereas prefusion SARS-CoV-2 spike can be observed in receptor-accessible conformations with one or more RBDs open^46,61,64–67^.

**Figure 4.**
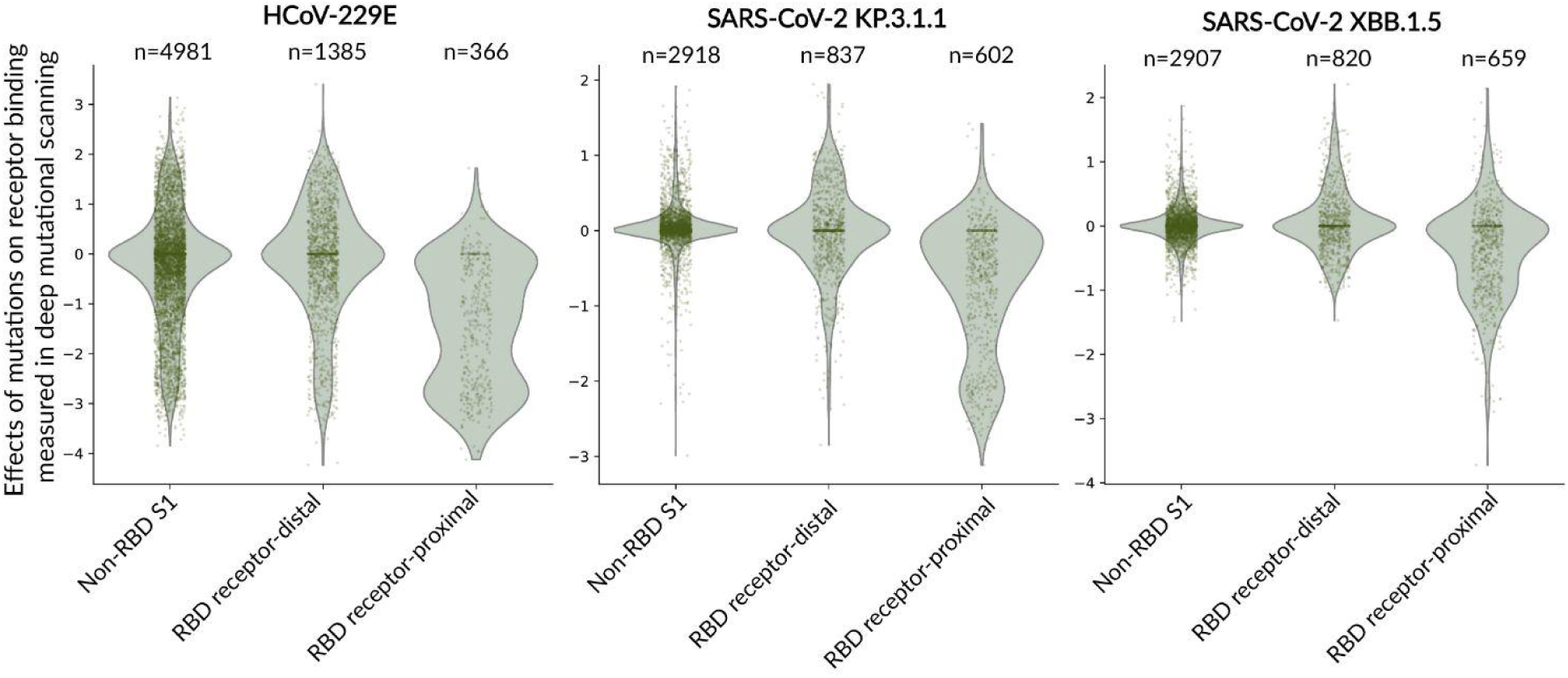
Effects of mutations in different regions of S_1_ on receptor binding as measured in the current study on the HCoV-229E spike and prior studies on the SARS-CoV-2 spike. Distribution of effects of mutations on receptor binding scores for all measured individual mutations for binding of the HCoV-229E spike to hAPN as measured in this study, and the binding of SARS-CoV-2 spikes from the KP.3.1.1 and XBB.1.5 spikes to human ACE2 as measured in previously published work, filtering only for mutations in the S_1_ subunit. Mutations are grouped by whether the site is outside the RBD, at a RBD site distal (>15 Å from) the receptor, or at a RBD site proximal (≤15 Å) to the receptor. Points represent individual mutations; violin plots show the smoothed distribution. The number of mutations in each group is indicated at the top of the plot. Note that the HCoV-229E deep mutational scanning covered nearly all spike mutations, the KP.3.1.1 deep mutational scanning covered all RBD mutations and non-RBD mutations that are observed in natural sequences or otherwise thought likely to be functionally tolerated, and the XBB.1.5 deep mutational scanning covered only mutations observed in natural sequences or otherwise thought to be functionally tolerated. See https://dms-vep.org/229E_spike_1984_DMS/htmls/coronavirus_comparison_interactive.html for an interactive version of this plot.

### Human sera that neutralize the HCoV-229E spike differ in their resistance to erosion by viral evolution

To understand how HCoV-229E spike evolution affects neutralization by human sera, we expanded prior experimental work by our group that tested the ability of historical human sera to neutralize pseudoviruses with spikes from both historical and more recent strains of HCoV-229E^5^. Specifically, we tested 24 human sera collected from adults between 1984 and 1995 for their ability to neutralize pseudoviruses with HCoV-229E spikes from 1984, 1992, 2001, 2008, and 2016 (Fig. S7); including 13 previously described sera^5^. The five virus strains for which these spikes are derived are the same used as in our prior work^5^ and are indicated in Fig. S1A and are detailed in Table S1; they differ by up to 44 amino-acid mutations in spike reflecting the rapid antigenic evolution of that protein^5,6^.

Concurring with our prior observations^5^, the neutralization of most of the historical human sera were eroded by subsequent viral evolution, with titers tending to be higher to older strains similar to ones the adults may have been exposed to prior to sera collection, and lower to later strains that evolved after the sera were collected (Fig. 5 and S7). However, among the sera there was clear variation in the extent of sensitivity to viral evolution: some sera had their neutralization rapidly eroded by viral evolution (Fig. 5, top row) whereas other sera continued to neutralize even spikes from well after the serum collection date albeit at lower levels (Fig. 5, bottom row). Based on these patterns, we classified five sera as “evolution-sensitive” and three sera as “evolution-resistant” as shown in Fig. 5.

**Figure 5.**
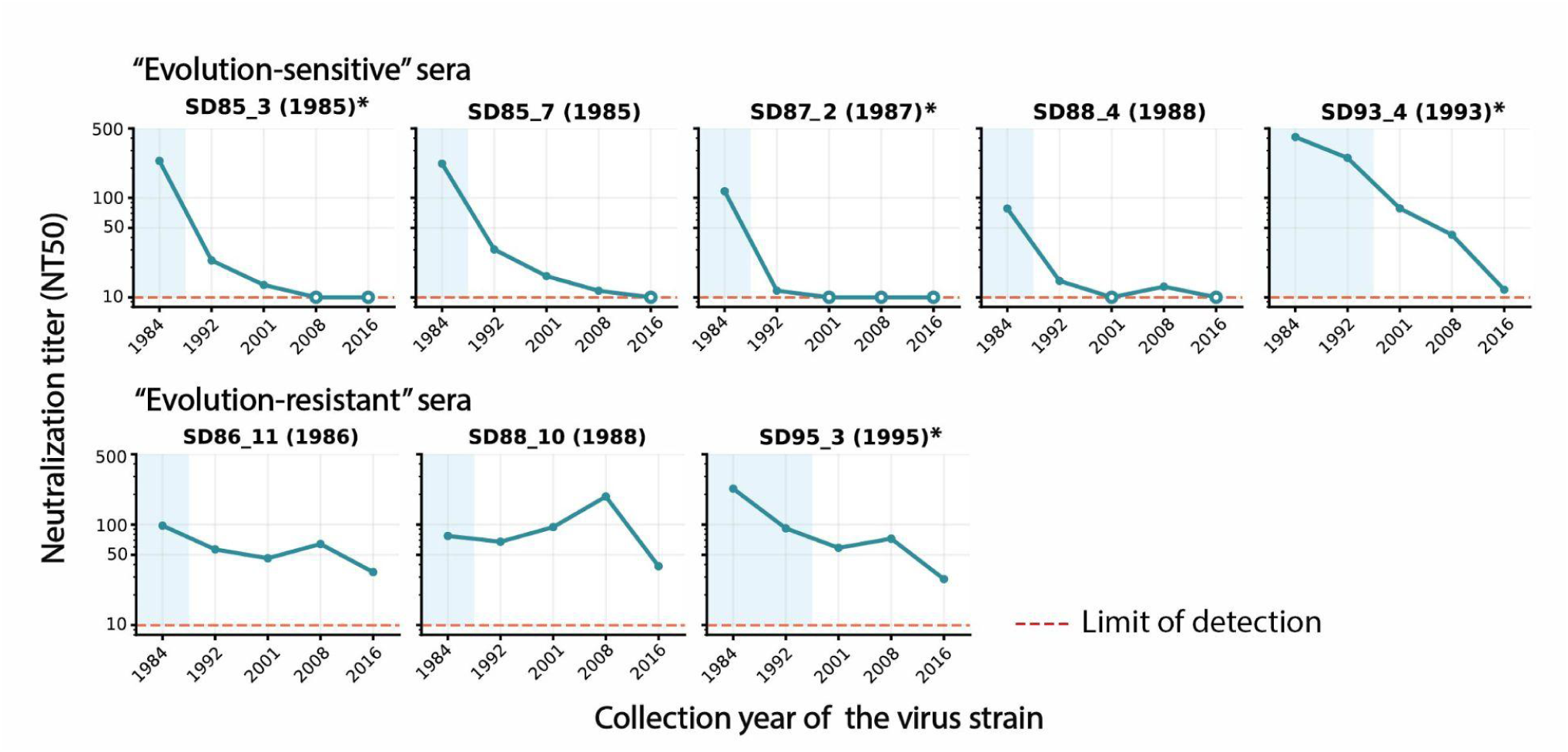
Neutralization of pseudoviruses with HCoV-229E spikes from different years by “evolution-sensitive” and “evolution-resistant” human sera. Neutralization by historical human sera of lentiviral particles pseudotyped with HCoV-229E spikes from strains collected in different years spanning 1984 to 2016. Titers are reported as the reciprocal of the serum dilution that neutralizes half the viral infectivity. Each facet shows neutralization of the pseudoviruses with spikes from different years by a distinct serum; the facet titles indicate the year the serum was collected and the blue shading indicates the interval prior to serum collection. Open symbols denote titers at the assay lower limit of detection of 10. These sera exemplify two patterns: the top row shows serum that are evolution-sensitive (neutralization rapidly eroded by viral evolution), and the bottom row shows sera that are evolution-resistant (retain the ability to neutralize viruses with spikes from many years after the serum was collected). Sera with an asterisk next to their name indicates that the neutralization data shown here has been previously published^5^.

### HCoV-229E spike mutations that affect RBD up/down conformation can be inferred from their effects on serum neutralization and hAPN binding

Previous studies of SARS-CoV-2 have shown that spike mutations that cause RBD up/down conformational changes have correlated effects on receptor binding and serum antibody neutralization, as mutations that promote the RBD up state increase receptor binding and susceptibility to neutralization, whereas mutations that promote the down conformation decrease both receptor binding and susceptibility to neutralization^53,54^. The reason for these correlated effects is that the conformational changes leading to the RBD opening expose the receptor-binding motif/loops as well as key RBD epitopes targeted by neutralizing antibodies.

We used deep mutational scanning to measure how spike mutations affect neutralization by all eight sera shown in Fig. 5. As schematized in Fig. S8, for these measurements we incubated the pseudovirus libraries across a range of serum concentrations, infected cells, and then extracted and sequenced viral barcodes, using a spike-in control to normalize barcode counts to calculate fraction viral infectivity as a function of serum concentration. We analyzed the data using a previously described framework^68^ to calculate the effect of each mutation on serum neutralization; these effects are reported as escape scores with positive values indicating a mutation reduces (escapes) neutralization and a negative value indicating a mutation increases serum neutralization.

The HCoV-229E spike mutations that most strongly affect neutralization are concentrated in the RBD and NTD (Fig. 6A-C and interactive figures at https://dms-vep.org/229E_spike_1984_DMS/RBD_up_down.html). The sites where mutations caused the greatest reduction in neutralization included 200, 210 and 213 in NTD, and 297, 309 and 336 in the RBD (Fig. 6A-C). There were more mutations that strongly increased neutralization (negative escape) than reduced neutralization (Fig. 6A-C). Most of these neutralization-sensitizing mutations are located in NTD (e.g., sites 90-93, 138, 141), although mutations at some RBD and S_2_ sites also increase neutralization (e.g., sites 326, 328, 431, 438, 658, 915, 924; Fig. 6B-C).

**Figure 6.**
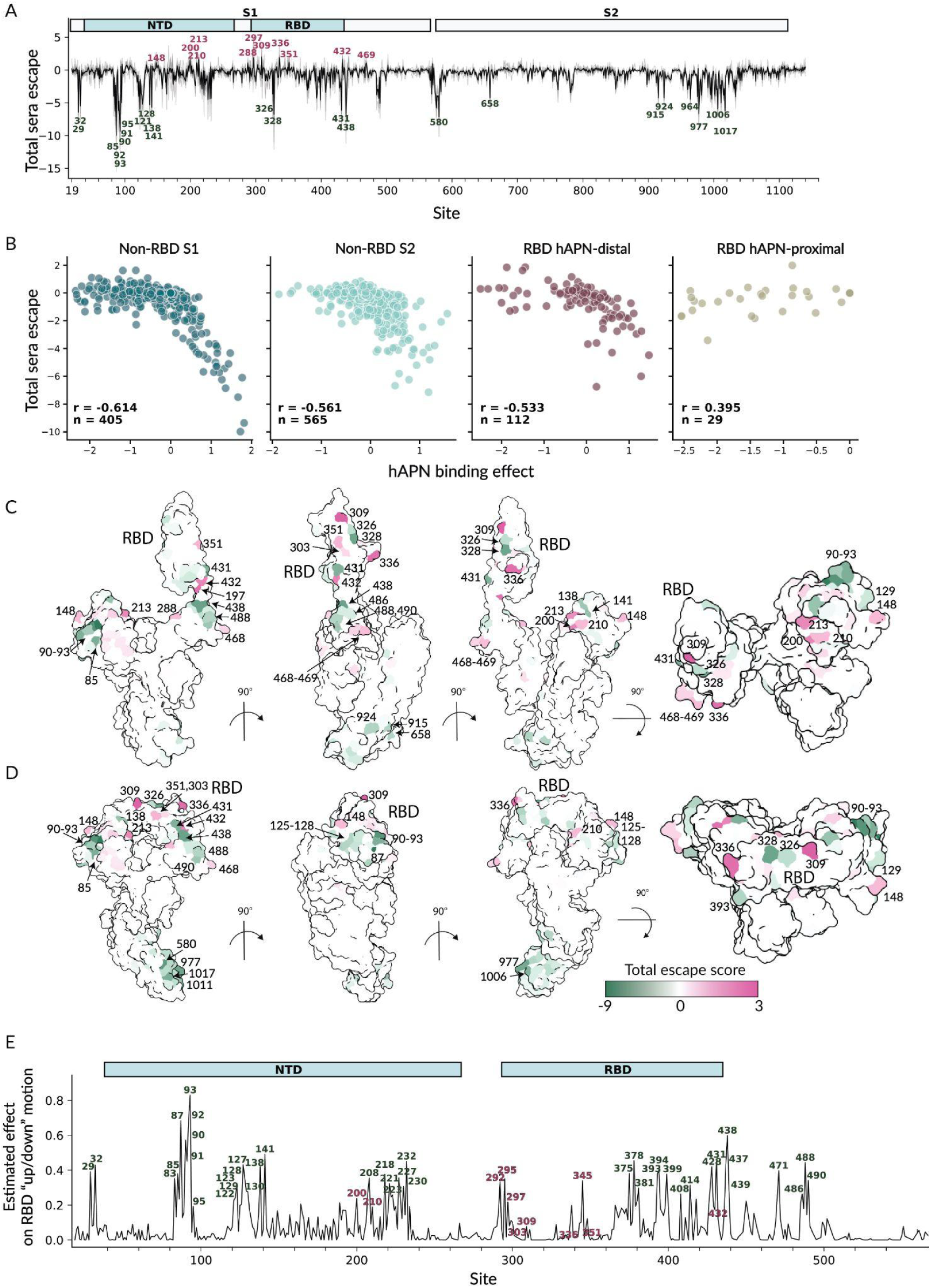
Effects of HCoV-229E spike mutations on serum neutralization and RBD up/down conformation. (A) Total effect of all functionally tolerated amino-acid mutations at each site on serum neutralization. Positive escape values indicate mutations reduce (escape) neutralization, while negative values indicate mutations increase neutralization. The thick black line shows the average over all eight sera in Fig. 5, while the thin gray lines show the values for individual sera. (B) Correlation between each mutation’s measured effect on hAPN binding and on serum neutralization, faceted by the site’s proximity to the hAPN binding motif. (C-D) Total serum escape per site averaged across all sera projected onto the spike structure with RBD up (panel C, PDB 8WDE) and RBD down (panel D, PDB 6H7U). Pink indicates sites where mutations reduce neutralization, and green indicates sites where mutations increase neutralization. (E) Estimated effect of S_1_ mutations at each site on RBD up/down conformation. Larger values indicate sites where mutations more strongly influence RBD up/down motion. Sites where mutations reduce serum neutralization (likely by putting the RBD more down) are labeled in pink, while sites where mutations increase serum neutralization (likely by putting the RBD more up) are labeled with green. See https://dms-vep.org/229E_spike_1984_DMS/RBD_up_down.html for interactive versions of the plots in this figure.

To assess the extent to which mutations might be affecting serum neutralization by modulating RBD up/down conformation, we examined the correlation between serum neutralization and hAPN binding (Fig. 6B). With the exception of the receptor-binding loops, there was a clear inverse correlation between mutation effects on escape from serum neutralization and receptor binding (Fig. 6B). This inverse correlation reflects the competing effects on these two properties caused by mutations that modulate RBD up-down conformation: mutations that promote RBD opening increase access to the receptor-binding motif for hAPN and also neutralizing antibodies. However, there is no such correlation for mutations mapping to the RBD proximal to the hAPN binding site, likely due to direct modulation of interactions with hAPN and neutralizing antibodies, compatible with most coronavirus polyclonal neutralizing activity target the RBD receptor-binding motif^25,48–51^. Although similar observations were made for the SARS-CoV-2 spike^53,54^, for the HCoV-229E spike mutations that strongly increase neutralization while increasing receptor binding are much more frequent and have greater effects. This difference likely reflects the fact that the HCoV-229E S trimer is more closed than the SARS-CoV-2 spike, meaning that there is more room for mutations to put the RBD in a more up conformation^37,47,62,63^.

To estimate how strongly each site affects RBD up/down motion, we applied a previously described approach^54^. Specifically, for each site we calculated the correlation between serum escape and hAPN binding effects across mutations at that site, and weighted this correlation by the root mean square of mutational effects on both phenotypes. Larger values indicate sites where mutations more strongly couple receptor binding and neutralization, indicating a greater impact on RBD conformation. Because this metric captures sites where mutations can promote either the up or down RBD state, sites with similar scores can reflect opposite directional shifts depending on the effects of individual mutations.

The sites in S_1_ estimated to have the greatest effect on RBD conformation are shown in Fig. 6E (see also the interactive plot at https://dms-vep.org/229E_spike_1984_DMS/RBD_up_down.html). Most mutations at sites that strongly impact up/down conformation increase serum neutralization (eg, are sensitizing rather than escape mutations), as indicated by the green labels in Fig. 6E. We do also identify a few sites (eg, 200, 210, and 297) where mutations affect RBD up/down conformation and reduce serum neutralization. However, the fact that most sites that affect RBD up/down conformation increase neutralization is consistent with the idea that the HCoV-229E spike is predominantly in the prefusion closed RBD state^37,47,62,63^, with most mutations that modulate conformation promote RBD opening thereby increasing serum neutralization.

### Effects of HCoV-229E spike mutations on neutralization by evolution-sensitive and evolution-resistant sera

We next sought to understand if there were differences in how spike mutations affected neutralization by the evolution-sensitive versus evolution-resistant human sera (as classified in Fig. 5). Many of the mutations that affect neutralization are the same across both groups of sera (Fig. 7A and interactive plots at https://dms-vep.org/229E_spike_1984_DMS/sera_escape.html). In particular, the mutations that increase neutralization (negative escape) are broadly similar for both groups of sera, suggesting that putting the RBD in a more up conformation increases neutralization by both groups.

**Figure 7.**
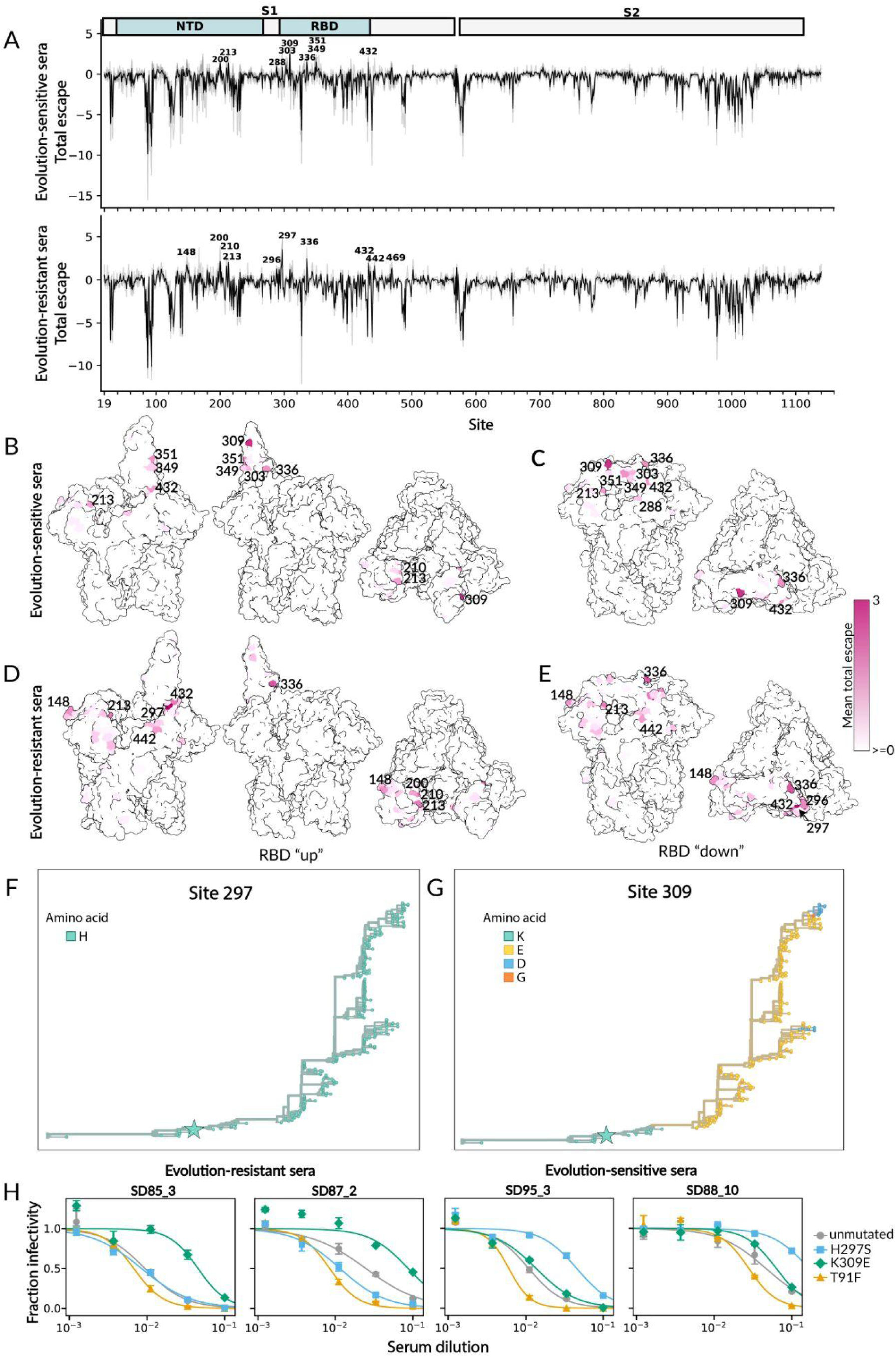
Effects of mutations to HCoV-229E spike on neutralization by evolution-sensitive and evolution resistant human sera. (A) Total effect of all functionally tolerated mutations at each site on neutralization by human sera, shown separately for the evolution-sensitive and evolution-resistant human sera from Fig. 5. The thin gray lines show individual sera, and the thick black line shows the average across the sera. See https://dms-vep.org/229E_spike_1984_DMS/sera_escape.html for interactive plots showing these data. (B-E) Total escape per site for the evolution-sensitive (B-C) and evolution-resistant (D-E) sera displayed on HCoV-229E spike structure with RBD up (PDB 8WDE; B and D) and RBD down (PDB 6H7U ; C and E). Dark pink indicates mutations reduce (escape) neutralization, while white means mutations do not impact or increase neutralization. (F-G) Amino-acid identity at spike sites 297 (F) and 309 (G) mapped onto the HCoV-229E phylogeny. Branches are colored by the inferred amino acid at the indicated site on that part of the phylogenetic tree. The star marks the 1984 strain used for deep mutational scanning. (H) Validation assays showing neutralization of lentiviral particles pseudotyped with HCoV-229E spike carrying the indicated mutation by two evolution-sensitive sera (SD85_3 and SD87_2) and two evolution-resistant sera (SD95_3 and SD88_10).

However, some of the mutations that cause most reduction (escape) in neutralization differ among the two sera groups (Fig. 7A-E). In particular, for the evolution-resistant sera most of the mutations that cause the greatest reduction in neutralization are at sites that the analyses in the preceding section suggest put the RBD in a more down conformation, such as 200 and 297. In contrast, the mutations that cause the greatest reduction in neutralization by the evolution-sensitive sera are mostly in or close to the RBD’s receptor binding loops and are accessible in both the up and down RBD conformation (eg, sites 303, 309, 349, and 351). This observation suggests that evolution-sensitive sera can erode neutralization by acquiring RBD mutations that directly reduce neutralizing antibody binding, whereas the neutralizing antibodies targeting in the evolution-resistant sera are escaped predominantly by putting the RBD in a more down conformation.

Mutations at sites that reduce neutralization by the evolution-sensitive sera are observed in the natural evolution of HCoV-229E (where as discussed above, the receptor-binding loops are the most variable region), but mutations that reduce neutralization by the evolution-resistant sera are generally not observed in natural evolution (Fig. 7F). This fact suggests it is often evolutionarily feasible for HCoV-229E to escape antibodies that directly target the receptor-binding loops via mutation in those loops, but mutations that put the RBD in a more down conformation might be too costly for other aspects of viral fitness even if they reduce serum neutralization.

To validate these deep mutational scanning results, we selected three mutations to individually test in pseudovirus neutralization assays: T91F, a neutralization-sensitizing mutation across all sera; H297S, an escape mutation for evolution-resistant sera; and K309E, an escape mutation for evolution-sensitive sera. We measured neutralization of each mutant using two evolution-resistant sera (SD88_10 and SD95_3) and two evolution-sensitive sera (SD85_3 and SD87_2). Consistent with the deep mutational scanning measurements, T91F increased serum neutralization across all sera (Fig. 7H). In contrast, H297S reduced neutralization by evolution-resistant sera, and K309E reduced neutralization by evolution-sensitive sera (Fig. 7H).

## Discussion

Here we used pseudovirus deep mutational scanning to understand how the HCoV-229E spike has evolved to erode antibody immunity while retaining its essential cell entry functions. We performed these experiments using the spike from a strain isolated in 1984, providing a historical reference point for comparing measured mutational effects to changes observed in later strains. We measured the effects of spike mutations on multiple phenotypes: cell entry into 293T cells expressing hAPN and TMPRSS2, binding to hAPN, and neutralization by human sera. The sera were pre-screened for their neutralization of a panel of historical and recent HCoV-229E spikes and grouped by whether their neutralization was relatively sensitive or resistant to viral evolution. This enabled us to assess the speed with which neutralizing activity is eroded by viral evolution across different human sera samples.

Our measurements of how mutations affect spike-mediated cell entry provide a comprehensive map of mutational tolerance, helping explain the rapid pattern of natural sequence change in HCoV-229E. During HCoV-229E spike’s evolution in humans the three hAPN-contacting RBD loops are the most variable region of spike ^5,52^, consistent with strong immune selection for mutations in and near the receptor-binding surface^5,6^, yet divergent strains maintain strong hAPN binding^52^. Our data explain this apparent disconnect by showing that most sites in the RBD’s receptor-binding loops are actually quite tolerant of mutations, and the handful of sites that have been conserved over decades of evolution are those where most mutations strongly impair spike-mediated cell entry. These results suggest that immune selection has driven mutational change at essentially all sites in the receptor-binding loops that can tolerate sequence change, with the only conserved sites being those where mutations are strongly incompatible with spike’s function.

We also directly measured how mutations affect spike’s binding to its hAPN receptor. A striking result is that some mutations at sites far from the RBD’s receptor-contact interface have large effects on receptor binding. This result can be understood in light of the facts that coronavirus RBDs can adopt both an up and down conformation, only the up RBD conformation can bind the receptor, and the HCoV-229E spike has evolved to strongly favor the RBD down conformation. Our results indicate that mutations outside the RBD’s receptor binding interface (especially in the NTD and receptor-distal regions of the RBD, but also in other regions of S_1_ and in S_2_), can have large apparent impact on receptor binding in the context of full spike by shifting the balance of up versus down RBD conformations. Notably, the mutations that promote receptor binding by putting the RBD in a more up conformation also increase neutralization by human serum antibodies, since they are better able to bind to key neutralizing epitopes that are exposed only in the RBD-up conformation. We speculate that long term evolution of HCoV-229E in humans has been selected for a strongly down RBD conformation to mask neutralizing epitopes in and near the RBD’s receptor-binding interface. Interestingly, a similar evolutionary pattern of selection for a more down RBD to reduce serum antibody neutralizing is becoming apparent in the evolution of SARS-CoV-2, with a variety of studies indicating that recent SARS-CoV-2 variants have more down RBDs than the earliest variants in the pandemic^54,61,69,70^.

Neutralization by historical human sera is generally weaker against spikes from more recent HCoV-229E strains, but the rate with which viral evolution erodes neutralization differs among sera from different individuals^5^. Our results suggest that this variation in the sensitivity or resistance of different sera to erosion by viral evolution reflect differences in the epitopes that are targeted. Evolution-sensitive sera are strongly affected by mutations in or near the RBD receptor-binding loops^5,6,25,37,52^, whereas evolution-resistant sera are more strongly affected by mutations that shift the RBD toward the down state and reduce epitope accessibility. Notably, mutations that reduce neutralization by evolution-resistant sera are uncommon in natural HCoV-229E evolution, suggesting that further masking RBD epitopes via the down conformation may have fitness costs (possibly reduced receptor binding) that are difficult for evolution to surmount.

## Limitations of study

Our study has several caveats. First, we performed the experiments in the background of a single HCoV-229E spike from a 1984 strain, which provides a useful historical reference point for interpreting long-term evolution but may not fully generalize to more recent strain backgrounds where epistasis could alter mutational effects. Second, all experiments were performed using pseudotyped lentiviral particles (pseudoviruses) in a standard cell line overexpressing the hAPN receptor. While this approach avoids the biosafety concerns related to mutagenizing authentic virus, pseudoviruses likely do not capture the full set of selective pressures acting on spike in the context of a replicating virus and natural infection.

## Acknowledgments

We thank Brendan Larsen for help with the library construction protocol and M. Alejandra Tortorici for her assistance with construct design. This study was supported by the NIAID/NIH under R01AI141707 (to JDB), P01AI67966 (to DV and JDB), DP1AI158186 and 75N93022C00036 (to DV) and an Investigators in the Pathogenesis of Infectious Disease Awards from the Burroughs Wellcome Fund (DV). JDB and DV are investigators of the Howard Hughes Medical Institute and DV is the Hans Neurath Endowed Chair in Biochemistry at the University of Washington. SH is a postdoctoral fellow of the Translational Data Science Integrated Research Center at the Fred Hutchinson Cancer Center. This research was also supported by the Genomics & Bioinformatics Shared Resource, RRID:SCR_022606, of the Fred Hutch/University of Washington Cancer Consortium (P30 CA015704), by the Flow Cytometry Shared Resource, RRID:SCR_022613, of the Fred Hutch/University of Washington/Seattle Children’s Cancer Consortium (P30 CA015704), and by Fred Hutch Scientific Computing, NIH grants S10-OD-020069 and S10-OD-028685.

## Material and methods

### Data and code availability

All data, interactive visualizations, and raw data from experiments described in this manuscript are publicly available on GitHub. The homepage at https://dms-vep.org/229E_spike_1984_DMS/ contains interactive visualizations to explore the deep mutational scanning data and links to additional datasets and associated raw data.

All code used to analyze the data and create figures is available on GitHub at https://github.com/dms-vep/229E_spike_1984_DMS.

### Biosafety

All experiments used spike-pseudotyped lentiviral particles (pseudoviruses), which encode no viral proteins other than spike are so only capable of undergoing a single round of cell entry, meaning that they are not fully replicative infectious agents capable of causing disease. No mutants of actual replicative virus were generated in this study.

### Sequencing numbering

Throughout the manuscript, spike residue numbering for the 1984 HCoV-229E spike in all deep mutational scanning data, follows RefSeq accession NC_002645.

### Deep mutational scanning library design for golden gate assembly

The spike gene from the HCoV-229E strain 1984 (NCBI accession DQ243964) was codon-optimized using the GeneArt tool (Thermo Fisher Scientific) and used as the wild-type template for deep mutational scanning (DMS) library construction. A full plasmid map containing the spike sequence and lentivirus genome used for this study is at https://github.com/dms-vep/229E_spike_1984_DMS/blob/main/data/paper_reference_files/plasmid_maps/4634_V5LP_pH2rU3_ForInd-Extgag_spike-229E-1984-opt2-d19_CMV_ZsGT2APurR.gb.

Libraries were generated using a previously described Golden Gate assembly strategy^54,71^, that in turn was based on prior work^72–78^, adapted for the 229E spike. To enable efficient mutagenesis and assembly, the spike coding sequence was divided into thirteen 297-bp tiles. Adjacent tiles were designed to overlap for two purposes: (i) to ensure complete mutational coverage of all spike coding sites across the tiled design, and (ii) to provide shared primer-binding regions that allowed each tile to be selectively PCR-amplified from a single pooled oligonucleotide library prior to assembly. For each tile, we synthesized an oligo pool encoding all 19 amino-acid substitutions (Twist BioSciences) at each position. In addition, we included premature stop-codon variants at 20 alternating positions within sites 20–40 (i.e., a stop at every other position) to serve as negative controls for the DMS cell-entry measurements. To allow golden gate assembly of the tiles, each PCR primer we used for tile amplification contains BsmBI cut sites. The full list of ordered tiles can be found here: https://github.com/dms-vep/229E_spike_1984_DMS/blob/main/data/paper_reference_files/GGA_library_files/229E-1984_spike_tails_opool.fasta.

### Library construction using Golden Gate assembly

To amplify each individual tile from a single ssDNA oligo pool, we performed 13 PCR reactions. For each reaction, we used Q5 High-Fidelity 2X Master Mix (New England Biolabs, Cat. No. M0492L), 0.5 µM of forward and reverse primer, and 1 ng of ssDNA oligo pool. Each reaction was started at 98°C for 30 s and then underwent 17 cycles of 98°C for 10 s, 60-69°C for 10 s, and 72°C for 25s. To amplify the corresponding flanking spike sequences for each tile, we used Q5 High-Fidelity 2X Master Mix, 0.5 µM of forward and reverse primer, and 1 ng of HCoV-229E spike coding lentiviral backbone. The PCR program varied between the different tile flaking regions, where annealing temperature and extension time differed between reactions. Denaturation step (98°C for 10 s), annealing time (30 s) and extension temperature (72 °C) were constant. Primer sequences and region-specific annealing temp and extension time for all PCR reactions are listed in the following table: https://github.com/dms-vep/229E_spike_1984_DMS/blob/main/data/paper_reference_files/GGA_library_files/PCR_rx_229E-1984_GGA_library.csv. Following PCR reaction, tiles and flaking spike fragments PCR were purified using Ampure XP beads (Backman Culture) using 0.8X ratio.

Mutagenized tiles were assembled together with their corresponding flanking spike fragments into the pGGAselect DNA shuttle vector (NEBridge Golden Gate Assembly Kit, BsmBI-v2; E1602L). Assembly reactions contained 100 fmol each of the amplified tile pool and matching flanking fragments, along with 50 fmol pGGAselect shuttle plasmid. Reactions were cycled 30 times between 42°C for 1 min and 16°C for 1 min, followed by a final incubation at 60°C for 15 min. Assemblies were purified using AMPure XP beads in a 1X ratio and eluted in 20 µL water. For transformation, 1 µL of purified assembly was electroporated into NEB 10-beta Electrocompetent E. coli (C3020K), recovered in 1 mL recovery medium at 37°C for 1 h with shaking, then pelleted and resuspended in chloramphenicol-containing LB for overnight outgrowth at 37°C with shaking to amplify the library. High transformation efficiency of ∼5×10^5^ to 1×10^6^ colonies per tile was estimated by plating serial dilutions on chloramphenicol agar and counting colony-forming units the next day. Each tiles shuttle plasmid libraries were then isolated using the QIAprep Spin Miniprep Kit (27106).

Next, each tile-specific shuttle-vector library was PCR-amplified using Q5 High-Fidelity 2X Master Mix, 10 ng plasmid library template, and 0.5 µM each of forward (5’-atg ttc gtt ctg ctg gtg gc) and reverse (5’- aga gcg tcg tgt agg gaa aga gtg tga tcc aac tag gcg cgc ctt atc acc gga tcg aag agg cg) primers. Cycling conditions were: 98°C for 30 s and then underwent 30 cycles of 98°C for 10 s, 60°C for 10 s, and 72°C for 1.5min, final extension of 72°C for 5 min was added. PCR products were gel purified to confirm recovery of the expected full-length spike fragment. Because spike is relatively large, we generated two pooled spike libraries: an S_1_ library comprising tiles 1–6 and an S_2_ library comprising tiles 7–13. Tiles were pooled at equal molarity, and the pooling was performed twice to create two independent replicates of each subunit library, yielding four libraries total (S_1_ replicate 1–2 and S_2_ replicate 1–2). Each pooled library was then PCR-amplified a second time for two purposes: (i) to add BsmBI recognition sites required for the subsequent Golden Gate assembly step, and (ii) to append a 16-nt unique barcode to each spike variant via an overhang on the reverse primer. These four libraries were amplified using Q5 High-Fidelity 2X Master Mix, 10 ng plasmid template, and 0.3 µM forward (5’- ggc tac cgt ctc gca cca tgt tcg ttc tgc tgg tgg c) and reverse(5’- ggc tac cgt ctc ata gaN NNN NNN NNN NNN NNN aga tcg gaa gag cgt cgt gta ggg aaa) primers, using the cycling conditions: 98°C for 30 s and then underwent 30 cycles of 98°C for 10 s, 60°C for 10 s, and 72°C for 1.5min, final extension of 72°C for 5 min was added..

Finally, each barcoded spike library was cloned into the lentiviral backbone using the NEBuilder Golden Gate Assembly Kit, as described above. The lentiviral backbone was based on the standard vector used previously in our lab^79,80^ but was modified for Golden Gate assembly by introducing a pair of BsmBI recognition sites flanking the spike insertion locus. Full plasmid map of lentivirus backbone used is available at https://github.com/dms-vep/229E_spike_1984_DMS/blob/main/data/paper_reference_files/plasmid_maps/4847_V5LP_pH2rU3_ForInd-Extgag_BsmBI_FlankingSites.gb. Plasmid map of the cloned HCoV-229E 1984 in the lentivirus backbone is available at https://github.com/dms-vep/229E_spike_1984_DMS/blob/main/data/paper_reference_files/plasmid_maps/4634_V5LP_pH2rU3_ForInd-Extgag_spike-229E-1984-opt2-d19_CMV_ZsGT2APurR.gb. To ensure that assembly occurred only at the intended junctions, any additional BsmBI sites elsewhere in the backbone were removed by synonymous sequence changes. Assembly products were electroporated into electrocompetent bacteria and amplified by liquid culture as described above. Electroporation efficiency was again verified, and we maintained at least 1×10^7^ colony-forming units per library replicate. Although spike libraries were made separately for S_1_ and S_2_, we subsequently combined the libraries for many of the experiments reported here. Cell entry measurements were conducted separately using the S_1_ and S_2_ libraries, and the results were subsequently combined computationally.

Human APN binding and sera selections measurements were performed using the combined full spike libraries.

### Production of cell-stored deep mutational scanning libraries

To generate the cell-stored deep mutational scanning libraries, we followed a previously described approach (Fig. S1C)^29^. Briefly, lentiviral backbones carrying the barcoded spike libraries were first used to produce VSV-G–pseudotyped lentiviral particles. Two 6-well plates of 293T cells were transfected with lentiviral helper plasmids (26_HDM_Hgpm2 (Addgene, 204152), 27_HDM_tat1b (Addgene, 204154), and 28_pRC_CMV_Rev1b (Addgene, 204153)) together with a VSV-G expression plasmid (Addgene, 204156). At 48h post-transfection, VSV-G–pseudotyped virus was harvested from the culture supernatant and used to transduce 293T-rtTA cells at a low multiplicity of infection (MOI < 0.01) to ensure that most cells received a single viral variant. Transduced cells were selected with puromycin, expanded, and cryopreserved as aliquots of >20 million cells in liquid nitrogen until use.

### Generation of VSV-G and spike protein expressing pseudovirus libraries

To produce pseudoviruses bearing spike proteins from the integrated cell libraries, 1.5 × 10^8^ 293T-rTTA cells were seeded into 5-layer flasks (Corning 353144) in tetracycline-free D10 medium supplemented with 0.1 μg/mL doxycycline to allow spike expression from the integrated genome. The next day, each flask was transfected with 50 μg each of the helper plasmids: 26_HDM_Hgpm2, 27_HDM_tat1b, and 28_pRC_CMV_Rev1b using BioT transfection reagent according to the manufacturer’s instructions. At 48 h post-transfection, supernatants were collected, filtered through a 0.45 μm SFCA Nalgene 500 mL Rapid-Flow filter unit (ThermoFisher 09-740-44B), and concentrated by ultracentrifugation (100,000 × g, 1 h, 4 °C). Viral pellets were resuspended in DMEM and stored at -80 °C for downstream deep mutational scanning experiments.

To generate VSV-G pseudoviruses from the integrated cell lines, 3 × 10^7^ cells were seeded into 15 cm dishes in tetracycline-free D10 medium. The following day, each dish was transfected with 7.5 μg each of the helper plasmids (26_HDM_Hgpm2, 27_HDM_tat1b, and 28_pRC_CMV_Rev1b) and 7.5 μg of 29_HDM_VSV_G encoding VSV-G. At 48 h post-transfection, supernatants were harvested, filtered through a 0.45 μm SFCA Nalgene Rapid-Flow filter unit, and concentrated using Lenti-X Concentrator (Takara 631232) according to the manufacturer’s instructions. Viral pellets were resuspended in D10. Concentrated VSV-G pseudoviruses were aliquoted and stored at -80 °C for a control in the cell-entry deep mutational scanning experiments.

Pseudovirus titers (transduction units, TU/mL) were quantified by flow cytometry as the fraction of ZsGreen-positive cells. Typical pre-concentration titers were ∼1 × 10^5^ TU/mL in 293T-hAPN-TMPRSS2 expressing cells, increasing to ∼5 × 10^6^ after concentration.

### Long-read PacBio sequencing to link mutations to barcodes

To generate the variant-barcode lookup table for the deep mutational scanning libraries, we produced VSV-G-pseudotyped viruses from the cell-stored libraries as described above. Importantly, this barcode linking was done after integration of the library into cells to account for recombination of the pseudodiploid lentiviral genomes prior to that stage. We used VSV-G pseudotyping at this stage to recover all variants from the cell pool, including those carrying spike mutations that are highly deleterious for spike-mediated entry. Approximately 10 million transduction units of the resulting VSV-G pseudoviruses were then used to infect 293T cells. Twelve hours post-infection, non-integrated viral genomes were recovered using the QIAprep Spin Miniprep Kit.

We performed two rounds of PCR to amplify the barcoded spike sequences from the recovered lentiviral genomes, using the minimum number of PCR cycles to limit strand switching. Amplicons were sequenced by long-read circular consensus sequencing on a PacBio Sequel IIe. For each barcode, we assigned a consensus variant sequence supported by at least two CCS reads. The variant–barcode lookup table for all HCov-229E libraries including combined libraries can be seen here https://github.com/dms-vep/229E_spike_1984_DMS/blob/main/results/variants/codon_variants.csv.

### Measurement of mutation effects on cell entry effect

To quantify the effects of mutations on cell entry, we followed the strategy described by Dadonaite et al^29^ Briefly, we infected parallel wells with (i) the pseudovirus library displaying different spike variants and (ii) a VSV-G pseudotyped library. The VSV-G library provides a baseline measure of library composition because VSV-G mediates entry independently of spike.

For each infection, 2 × 10^6^ of 293T-hAPN-TMPRSS2 and 293T cells in D10 medium, were seeded per well of a 6-well plate. The next day, cells in two replicate wells were infected with 3 × 10^6^ TU of the spike-pseudotyped library or 10 × 10^6 TU of the VSV-G library. At 12 h post-infection, non-integrated reverse-transcribed lentiviral genomes were recovered by miniprep.

Illumina sequencing libraries were generated in two PCR steps. In Round 1, TruSeq Read 1 and Read 2 sequences were appended. Each 50 µL reaction contained 30 µL miniprepped DNA, 25 µL KOD Hot Start Master Mix, 1.5 µL of 10µM Illumina_Rnd1_For primer (5’- CTC TTT CCC TAC ACG ACG CTC TTC CGA TCT), and 1.5 µL of µM Illumina_Rnd1_Rev3 primer (5’- CTG GAG TTC AGA CGT GTG CTC TTC CGA TCT gtc cct att ggc gtt act atg gga aca tac gtc). Cycling was: 95 °C for 2 min; then 27 cycles of 95 °C for 20 s, 70 °C for 1 s, 58 °C for 10 s (0.5 °C/s ramp), and 70 °C for 20 s; followed by 70 °C for 60 s and a 4 °C hold. Round 1 products were purified with 150 µL AMPure XP beads and eluted in 50 µL nuclease-free water, and DNA was quantified on a Qubit 4 Fluorometer (ThermoFisher Q33238).

Next, Round 2 PCR was performed to add sample indices for multiplexing. In brief, each 50 µL reaction contained 20 ng of Round 1 product, 25 µL KOD Hot Start Master Mix, 2 µL each indexing primer (10 µM), and nuclease-free water to volume. Cycling conditions matched Round 1 except that 20 cycles were performed. Round 2 products were quantified by Qubit, pooled equimolarly, run on a 1% agarose gel, and the ∼300 bp band was excised. DNA was extracted using the NucleoSpin Gel and PCR Clean-up Kit and further purified with AMPure XP beads. Final libraries were diluted to 10 nM and sequenced on an Illumina NovaSeq X Plus.

Mutation effects on cell entry were quantified as a log2 enrichment ratio based on this equation:

*log_2_([counts of variant in spike-pseudotyped libraries / counts of variant in VSV-G-pseudotyped libraries] / [counts of all wildtype variants in spike-pseudotyped libraries / counts of all wildtype variants in VSV-G-pseudotyped libraries])*

where *counts of variant in spike-pseudotyped libraries* is the count of a variant after infection and the *counts of variant in VSV-G-pseudotyped libraries* is the count of the same variant in the pre-infection baseline measured using the VSV-G pseudotyped library, and *counts of all wildtype variants in s[pike-pseudotyped libraries*, *counts of all wildtype variants in VSV-G-pseudotyped libraries* are the corresponding counts for the unmutated (wild-type) variant. To infer single-mutation effects from libraries containing both singly and multiply mutated variants, we used the multidms packag^36^ to fit global epistasis model^35^ to the variant-level data. To ensure high quality data, we required each mutation to be observed in at least two unique barcodes (*times_seen >= 2*) and retained only mutations with low variability across replicate measurements (*effect_std <= 2.5*). Interactive plots containing the per mutation effects on cell entry are available at https://dms-vep.org/229E_spike_1984_DMS/htmls/cell_entry_func_effects.html

### Production and purification of soluble human APN

In order to produce soluble human APN, Expi293 cells (ThermoFisher Scientific) were cultured at 37°C in a humidified 8% CO_2_ incubator with constant rotation and transfected using the ExpiFectamine Transfection Kit (ThermoFisher Scientific) according to the manufacturers guidelines and recommendations. The transfected cells were cultured for five days and the supernatant clarified using centrifugation before being run over a HiTrap™ Protein A HP column (Cytiva). The columns were washed with 20 mM Sodium Phosphate pH 8.0 before the protein was eluted using 0.1 M citric acid pH 3.0 into 1.0 M Tris pH 9.0 to a final eluate pH of 8.0. The hAPN was then concentrated using Amicon Ultra-15 Centrifugal Filter Units 30kDa (Millipore) and run through a Superose 6 Increase column (Cytiva) equilibrated in 25 mM Tris, 150 mM NaCl pH 8.0. The fractions containing hAPN were pooled and flash frozen in liquid nitrogen and stored at -80°C.

### Measurement of mutation effects on receptor binding

To quantify how spike mutations affect hAPN binding, we used a soluble dimeric hAPN neutralization assay. For each sample, we combined 1.5 × 10^6^ TU of the spike-pseudotyped library with VSV-G-pseudotyped virus at 2% of total transcription units. VSV-G psuedotyped lentivirus production and use were described previously^29,81^. VSV-G serves as a non-neutralizable internal standard that enables conversion of sequencing counts to the fractional neutralization of each variant at each hAPN concentration. We then incubated the library with increasing concentrations of soluble dimeric hAPN at 37°C for 1h. hAPN concentrations were chosen to span the neutralization range of HCoV-229E spike-pseudotyped virus and thereby capture mutations that increase hAPN binding (variants neutralized at lower HAPN concentrations) as well as mutations that decrease binding (variants requiring higher hAPN concentrations for neutralization). The concentrations used were 5, 14, 35, 63, and 127 µg/mL. Following incubation, the libraries were used to infect 293T-hAPN-TMPRSS2 cells.

Non-integrated viral genomes were recovered and prepared for Illumina sequencing as described in the “Measurement of mutation effects on cell entry effect” section. Sequencing counts were converted to fractional neutralization using the VSV-G standard, and mutation effects were inferred using a biophysical model implemented in the polyclonal software package (https://github.com/jbloomlab/polyclonal^68^). We report effects such that positive values indicate improved hAPN binding (i.e., greater neutralization by soluble hAPN). Experiments were performed for each S_1_ and S_2_ libraries (1 and 2) biological replicates, and the values reported are the mean across replicates. To reduce experimental noise, we require each mutation to be observed in at least two unique barcodes (*times_seen >= 2*) and we only retain mutations with low variability across replicate measurements (*binding_std => 1.8*). In addition, to avoid poor measurements affected by impaired cell entry, we filtered out mutations with cell entry effect lower than -2.5. An interactive plot showing the per mutation effects on hAPN binding is available at https://dms-vep.org/229E_spike_1984_DMS/htmls/human_APN_binding_mut_effect.html.

### Measurement of mutation effects on serum neutralization

Before initiating serum selection experiments with the deep mutational scanning libraries, we first quantified the potency of each sample using pseudovirus neutralization assays against viruses pseudotyped with the HCoV-229E 1984 spike. Neutralization assays were performed as previously described^82^ and the mutant spike pseudovirus was generated as described in “<u>Production of</u> <u>luciferase-encoding pseudovirus for all validation assays</u>” section below. Fraction infectivity at each dilution was determined relative to serum-free controls, and the neutcurve package^83^. All sera were heat-inactivated at 56°C for 1 h prior to use.

For each selection, 1.5 × 10^6^ TU of the spike-pseudotyped library virus were mixed with increasing concentration of serum, and included a no-serum control. The starting serum dilution was set to 3× the IC99 measured in the standard pseudovirus neutralization assay; this standard assay typically underestimates neutralization in the deep mutational scanning selections, potentially due to differences in spike density between standard pseudoviruses and the library virus, and/or depletion of antibody molecules by the higher virion concentrations used in library experiments. After 1h incubation of spike-pseudotyped library virus and serum sample in 37°C, the mixtures were used to infect 293T-hAPN-TMPRSS2 cells At 12 h post-infection, non-integrated viral genomes were recovered and processed for Illumina sequencing as described “Measurement of mutation effects on cell entry effect” section.

To infer mutations affecting serum or antibody neutralization, we used the biophysical model implemented in the polyclonal package, as deployed in dms-vep-pipeline-3 (v3.29.0; https://github.com/dms-vep/dms-vep-pipeline-3). The serum neutralization data were filtered using three different cutoffs: mutations occur with at least two unique barcodes, minimum cell entry effect of -2.5 and standard deviations between replicates <= 2. Per sample interactive serum escape plots, are available at https://dms-vep.org/229E_spike_1984_DMS/sera_escape.html.

### Production of luciferase-encoding pseudovirus for all validation assays

Desired mutations were introduced into the HCoV-229E 1984 spike expression plasmid and plasmid sequences were verified by whole-plasmid sequencing. Spike-pseudotyped lentiviruses were generated by transfecting 293T cells with the spike expression plasmids together with a lentivirus helper plasmids (26_HDM_Hgpm2 (Addgene, 204152), 27_HDM_tat1b (Addgene, 204154), 28_pRC_CMV_Rev1b (Addgene, 204153)) and the pHAGE6_Luciferase_IRES_ZsGreen lentiviral backbone. Virus-containing supernatants were harvested 48 h post-transfection and filtered. To titrate these pseudoviruses, we seeded 60,000 293T-hAPN-TMPRSS2 per well in black-wall, clear-bottom, poly-l-Lysine coated 96-well plates 24 hours prior to infections. We used six 5-fold serial dilutions, starting dilution of 0.05. 48 hours post infection, we measured RLUs using the Bright-Glo Luciferase Assay System (Promega, Ref. No. E2620). For each virus, we calculated an RLU/μL from technical replicates by taking the mean of all individual replicates, and for each mutation we titrated two biological replicate viruses.

### Neutralization curves generation for all tested human sera

HCoV-229E spike proteins of different HCoV-229E strains (that were used in this study) were codon optimized and cloned into expression plasmids. Pseudotyped lentivirus particle was generated using the same protocol described in the method section “Production of luciferase-encoding pseudovirus for all validation assays”. Plasmid maps of the spike expression plasmids that were used can be found here https://github.com/dms-vep/229E_spike_1984_DMS/tree/main/data/paper_reference_files/plasmid_maps. Neutralization assay of all tested spikes were generated as previously described ^5^. In brief, 293T cells were transiently transfected to express hAPN and TMPRSS2 using expression plasmids using Bioland Scientific BioT transfection reagent following the manufacturer’s protocol. 6 hours post transfection, cells were seeded in 96-well-plate, and neutralization assays were performed 20–24 hours. Heat-inactivated sera were first diluted 1:10 in D10 growth medium and then subjected to 3-fold serial dilutions in TC-treated 96-well “set-up” plates to generate seven total dilutions per sample. Each dilution series was prepared in duplicate.

Each 229E spike-pseudotyped lentivirus was diluted to yield luciferase signals of ∼200,000 RLUs per well. Equal volumes of diluted virus were added to each serum dilution in the set-up plates and incubated for 1 hour at 37°C. After incubation, 100 µL of each virus-serum mixture was transferred onto the transfected 293T expressing hAPN and TMPRSS2 cell plates seeded the previous day. Plates were incubated at 37°C for ∼50–52 hours, after which luciferase was measured as in the titration assays.

Fraction infectivity at each dilution was calculated as the signal relative to the no serum control (averaged across the two control wells for each row) after subtracting the background calculated from cells only wells and using the neutcurve package^83^.

### Estimate of mutation effects on RBD up/down motion

We measured each site’s effect on RBD up/down motion using a similar approach to one published previously^54^. We used the following formula:

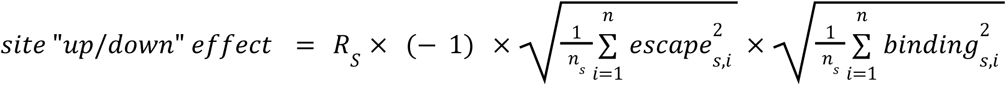

where *R* is the Pearson correlation, for site *s*, between mutation effects on serum escape averaged across all sera and hAPN binding. *R* positive values were set to 0, and the remaining values were multiplied by −1.

The root-mean-square (RMS) serum-escape effect at site s was computed as:

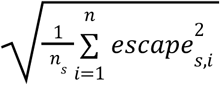

where *escape_s,i_* is the serum-escape effect of mutation *i* at site *s* (averaged across all sera) and *n_s_* is the number of mutations measured at site *s*. The RMS effect on hAPN binding was calculated analogously as:

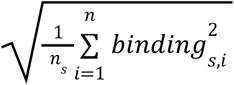

To ensure reliable inference of RBD up/down motion, we excluded mutations with extremely poor cell entry, therefore applied the same filter listed in the methods section “Measurement of mutation effects on cell entry effect “, and included mutations with cell-entry effects >= -2.5. To increase the analysis confidence, we display sites with measurements for at least five mutations.

### Phylogenetic tree generation

We gathered publicly available HCoV-229E spike sequences from NCBI Virus using the query term “Human Coronavirus 229E, taxid: 11137” in the “Search by virus name or taxonomy box.” Sequences were downloaded as of Jan 9th, 2026. Full list of NCBI accessions identifiers of the sequences that were used in the phylogenetic tree is available here https://github.com/jbloomlab/cov-229E-spike-phylo/blob/main/Results/spike/ORF_sequences/accession_numbers_after_filter.txt. Only spike sequences with at least 3400 nucleotides and no ambiguous nucleotides (“N”) were included. In addition, sequences that were shown as extreme outliers on date-to-tip regression were also filtered out. The spike sequences were aligned using Augur^84^, and a phylogenetic tree was inferred from the resulting alignment. The phylogenetic tree was rooted using Camel alphacoronavirus (NCBI accession: NC_028752) as an outgroup. Then, the outgroup was removed and we produced a timetree using TreeTime^85^. The tree was visualized using Nextstrain^86^.

See https://nextstrain.org/community/jbloomlab/cov-229E-spike-phylo@main for the interactive Nextstrain trees, and https://github.com/jbloomlab/cov-229E-spike-phylo for the computer code used to generate the tree.

### Natural variation analysis of the spike protein and the RBD loops

For natural variation analysis, we used the same alignment as for the phylogenetic analysis, and can be found here https://github.com/jbloomlab/cov-229E-spike-phylo/blob/main/Results/spike/Alignments/protein_ungapped_no_outgroup.fasta. Amino-acid diversity across HCoV-229E spike was quantified per site using the effective number of amino acids (2^*H*^), where *H* is Shannon entropy, so values range from 1 (fully conserved) upward as diversity increases. Diversity was calculated for 1,173 spike sites.

For the RBD loop comparison, we compared several selected HCoV-229E spike protein sequences (NCBI accession numbers from the top to bottom of Fig. 2B: AAK32191.1, DQ243964.1, ABB90507.1, DQ243976.1, DQ243986.1, KY369909.2, ON791801.1), all sequences were aligned based on the reference strain AAK32191.1.

### Structural visualizations

Structural figures were generated in UCSF ChimeraX (v1.10.190) using the Protein Data Bank accession IDs specified in the figure legends. ChimeraX filtered files with summarized deep mutational scanning data can be found here https://github.com/dms-vep/229E_spike_1984_DMS/tree/main/results/structure_analysis_plots/chimerax/results.

## Supplementary Figures

**Supplementary Figure S1.**
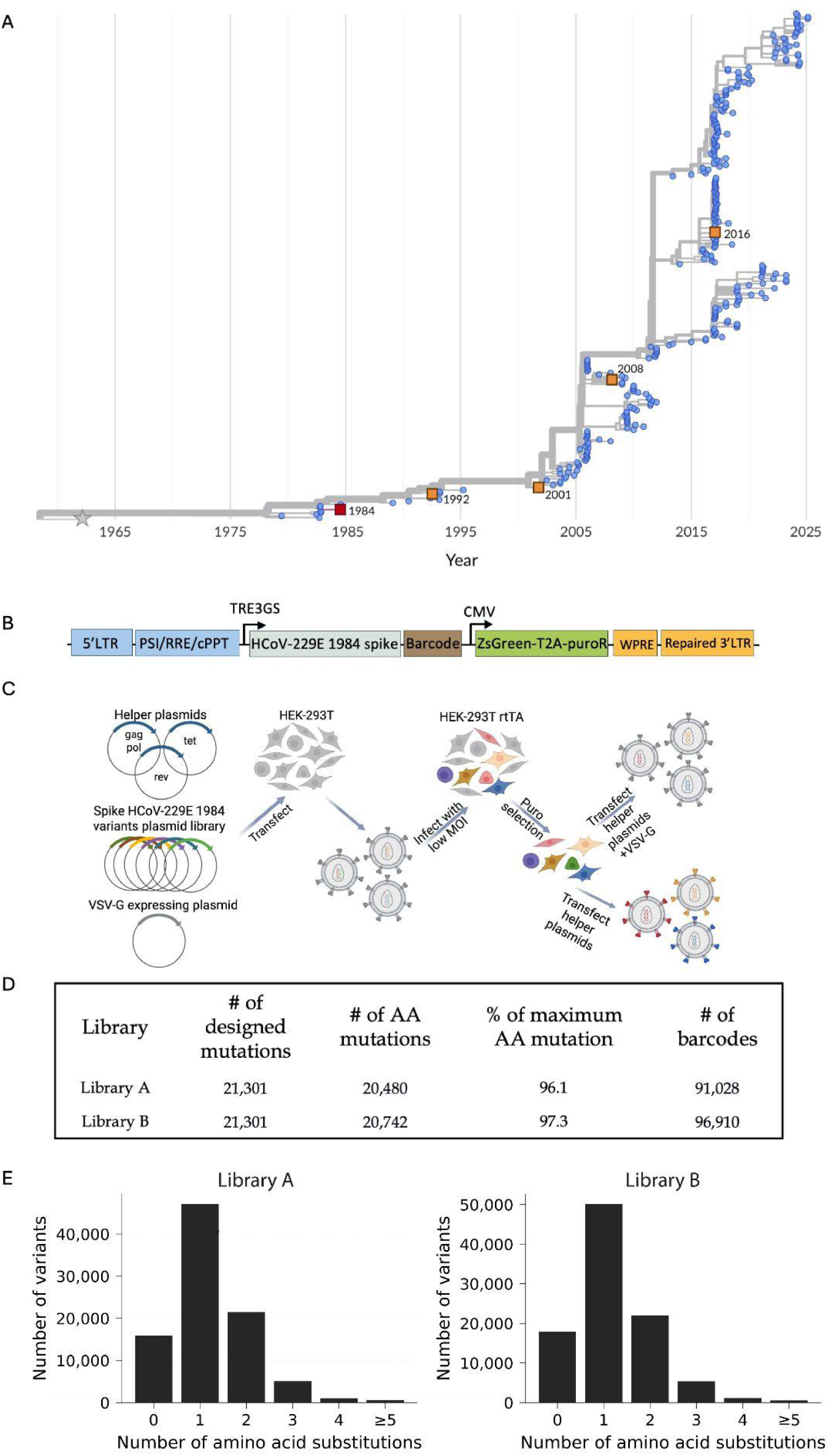
Pseudotyped lentivirus spike libraries design and production. (A) Time-scaled phylogeny of the HCoV-229E spike inferred using the full spike nucleotide sequences. Red square marks the HCoV-229E 1984 spike that is used for this study. Additional strains used in neutralization assays to identify evolution-sensitive versus evolution-resistant sera are marked with orange squares. The reference strain (NC_002645) from a 1962 isolate, marked as grey star, was extensively passaged in the lab prior to sequencing. See https://nextstrain.org/community/jbloomlab/cov-229E-spike-phylo@main for an interactive version of this phylogenetic tree. (B) Schematic of the lentiviral genome (backbone) used for pseudovirus deep mutational scanning^143^. The backbone contains standard lentiviral elements (5′ LTR, packaging signal Ψ, RRE, cPPT, and 3′ LTR), with a full-length (non-deleted) 3′ LTR to enable reactivation of integrated proviruses. It encodes the HCoV-229E spike protein under an inducible TRE3GS promoter, followed by a random nucleotide barcode placed downstream of the stop codon. A constitutive CMV promoter drives expression of ZsGreen and a puromycin-resistance cassette. (C) Workflow to generate a library of pseudotyped lentiviral particles. 293T cells are transfected with the library of plasmids encoding the spike variants together with lentiviral helper plasmids (encoding Tat, Gag-Pol, and Rev), and a plasmid expressing VSV-G. The resulting virions are used to infect 293T cells expressing rTTA at low multiplicity of infection (0.01). Infected cells with an integrated lentiviral genome are selected using puromycin; due to the low multiplicity of infection nearly all of these cells contain just a single integrated genome encoding a barcoded spike variant. The genotype-phenotype linked lentiviral library is then produced by retransfecting these cells with the helper plasmids. To quantify library composition independent of the spike function, particles pseudotyped with VSV-G are also generated and used as a control to normalize library composition independent of spike function. This panel was created with BioRender.com (D) Number of designed (intended) unique amino-acid mutations to be covered in each library, actual number of unique mutations represented at least once in each library, and number of uniquely barcoded spike variants in each library. (E) Distribution of the number of amino-acid mutations per barcoded variant for each library.

**Supplementary Figure S2.**
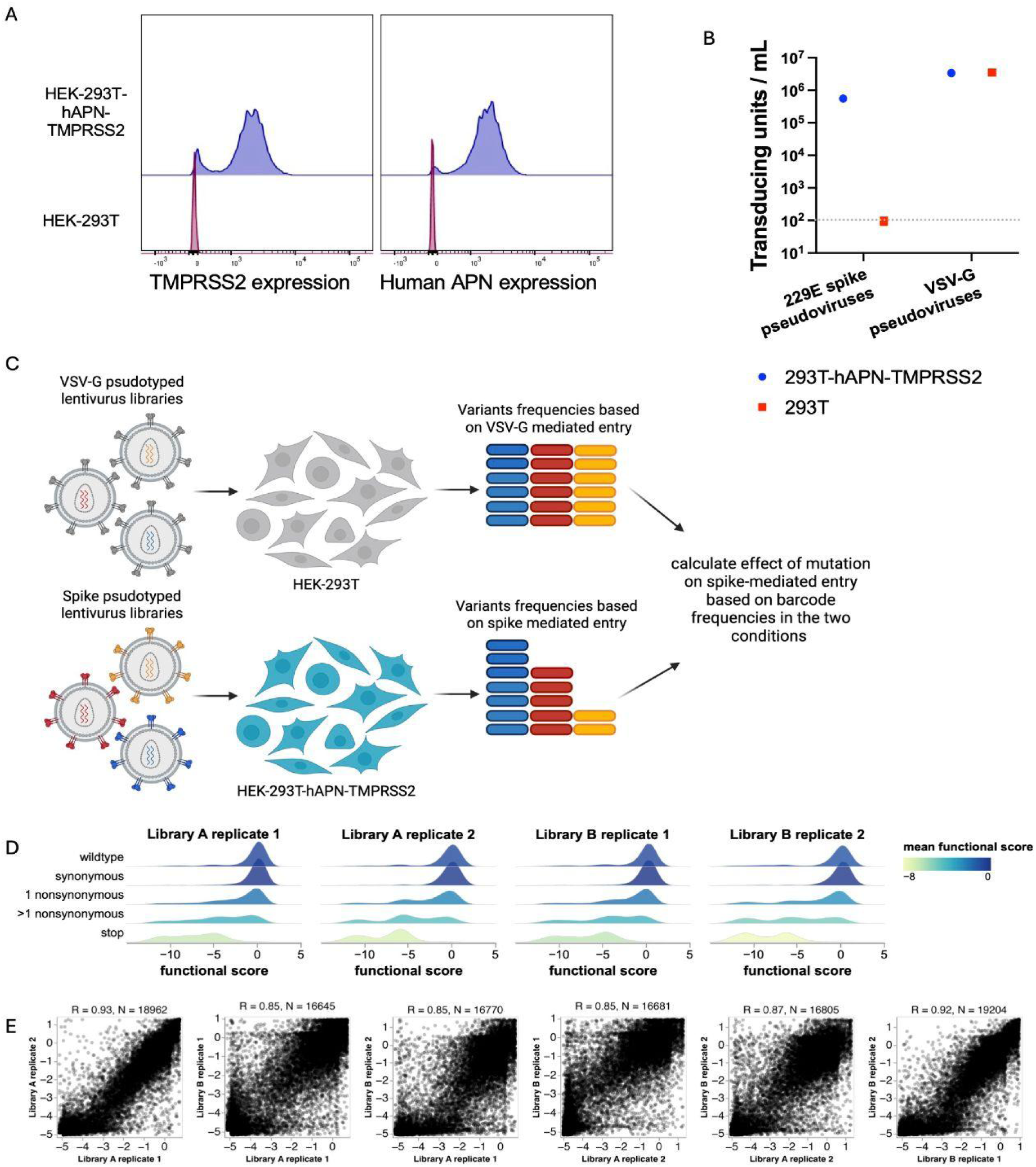
Measurement of effects of spike mutations on cell entry. (A) Expression of human amino-peptidase N (hAPN) and TMPRSS2 in the 293T-hAPN-TMPRSS2 cell clone used in this paper. Expression of hAPN was quantified by antibody staining and flow cytometry, while expression of TMPRSS2 was quantified by flow cytometry analysis of fluorescence of a mCherry reporter expressed off the same transcript. (B) Infectious titers of lentiviral particles pseudotyped with either the 229E spike or VSV-G, measured on 293T and 293T-hAPN-TMPRSS2 target cells. As expected, spike pseudotyped viral particles can only efficiently infect the cells that express the hAPN receptor and TMPRSS2 activating protease. (C) Schematic of deep mutational scanning workflow for measuring entry into 293T-hAPN-TMPRSS2 cells. Cells are infected with either the VSV-G pseudotyped library or the spike pseudotyped variant library produced as described in Fig S1. At 12 hours post-infection, viral DNA is extracted from infected cells, and barcode frequencies are quantified by sequencing^143^. All variants can enter cells equally when pseudotyped with VSV-G, but for the spike-pseudotyped virions entry is dependent on spike function. A cell entry score is computed by comparing barcode frequencies between the spike library and the VSV-G control library conditions. This panel was created with BioRender.com (D) Distributions of cell entry scores for variants grouped by the type of mutation they contain. A score of zero corresponds to entry equivalent to the unmutated spike, while negative values indicate reduced entry. (E) Correlation of per-mutation cell entry effects measured in two technical replicate experiments across each of the two independent biological replicate libraries, after inferring mutation-level effects from variant scores using global epistasis models (Methods). Numbers above plots give the Pearson correlation (R) and number of mutations (N) measured in both pairs of replicates.

**Supplementary Figure S3.**
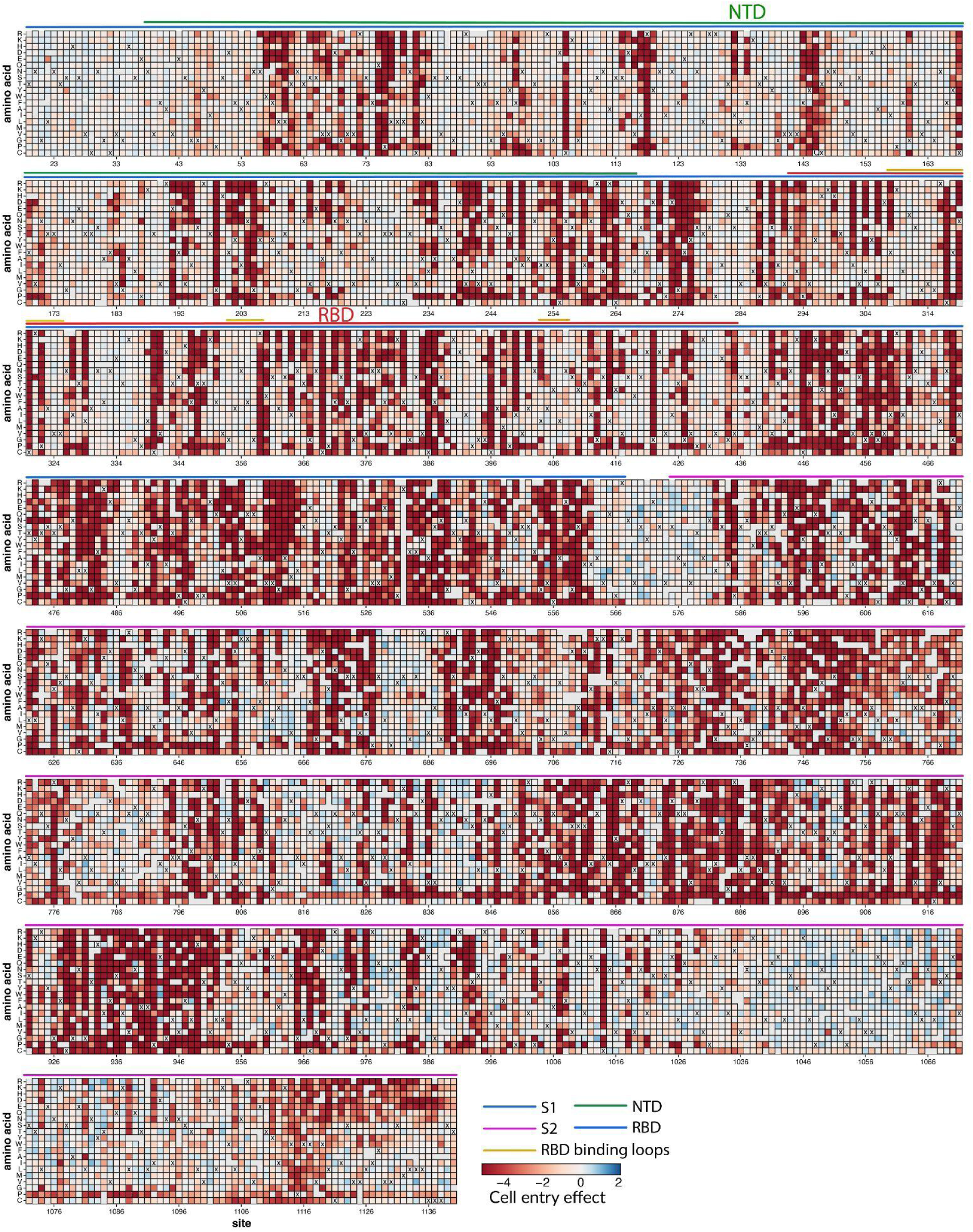
Effects of mutations to the HCoV-229E spike on entry in 293T-hAPN-TMPRSS2 cells. Each square in the heatmap represents the effect of an amino-acid mutation, with red indicating impaired entry, white indicating wildtype-like entry, and blue indicating improved entry. The wildtype amino acid of the 1984 HCoV-229E spike used for the deep mutational scanning is indicated with an “X” at each site. The handful of gray squares indicate mutations that were not measured with high confidence in the deep mutational scanning. The lines above the heatmap indicate different regions of spike. See https://dms-vep.org/229E_spike_1984_DMS/cell_entry.html for an interactive version of this heatmap.

**Supplementary Figure S4.**
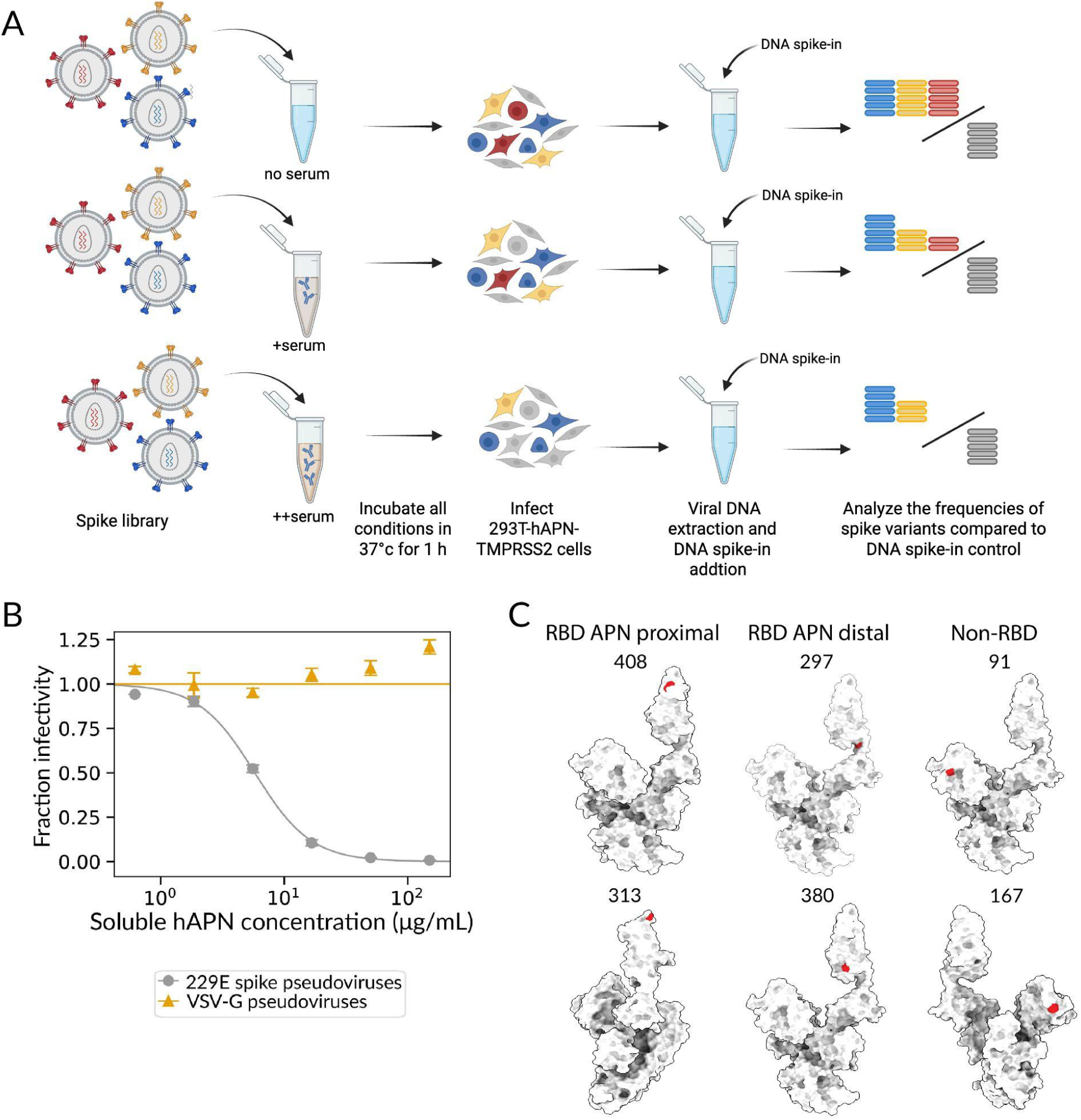
Measurement of effects of spike mutations on hAPN binding. (A) Schematic of deep mutational scanning to measure the effects of spike mutations on hAPN binding using 293T-hAPN-TMPRSS2 cells. The spike pseudotyped variant library is mixed with barcoded VSV-G pseudotyped lentiviral particles that act as a standard for normalization of sequencing counts, and incubated with increasing concentrations of soluble hAPN, including a no hAPN control. After 1h incubation at 37C°, cells are infected with the pseudovirus mix, and at 12 hours post-infection the viral DNA is extracted from infected cells, and barcode frequencies are quantified by sequencing^29^. VSV-G barcode counts are used to normalize the read counts of spike variants across conditions to determine the absolute infectivity of each spike variant at each hAPN concentration. Mutations that decrease receptor binding are neutralized less potently by soluble hAPN, while mutations that increase binding are neutralized more potently by soluble hAPN. This panel was created with <u>BioRender.com</u>. (B) Neutralization of pseudoviruses expressing the unmutated spike of HCoV-229E pseudoviruses or VSV-G by soluble hAPN. (C) Sites with mutations that were chosen for additional validations, displayed on one monomer of the spike protein structure PDB 8WDE^47^, segregated based on their proximity to the hAPN binding as defined in Fig 3.

**Supplementary Figure S5.**
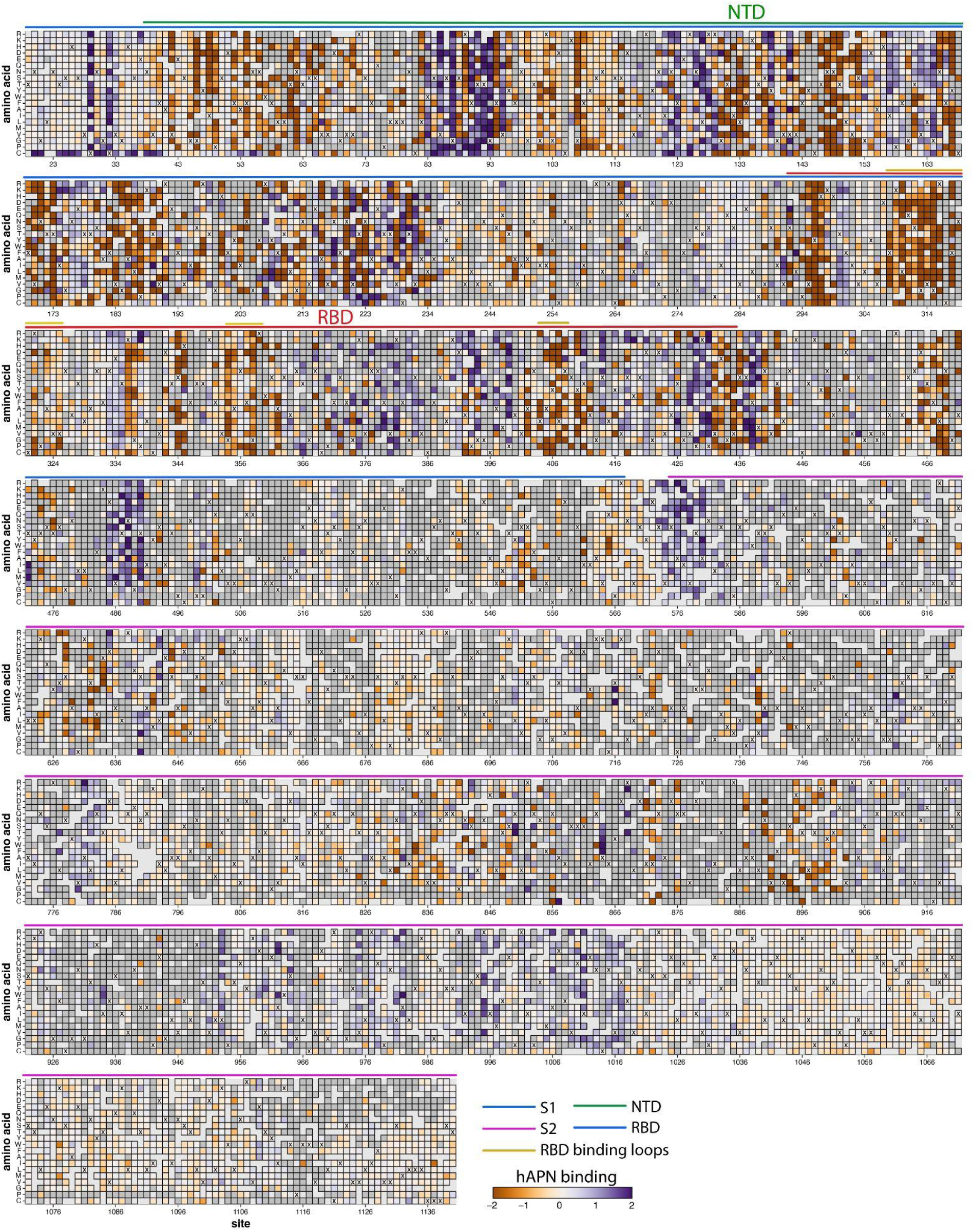
Effects of mutations to the HCoV-229E spike on hAPN binding as measured by pseudovirus neutralization by soluble hAPN. Each square in the heatmap represents the effect of an amino-acid mutation, with orange indicating reduced hAPN binding, white indicating wildtype-like hAPN binding, and purple indicating improved hAPN binding. The wildtype amino acid of the 1984 HCoV-229E spike used for the deep mutational scanning is indicated with an “X” at each site. Dark gray squares are mutations that strongly impair cell entry (*cell entry effect <=−2.5*), and therefore cannot be reliably measured for their impact on receptor binding. The handful of light gray squares indicate mutations that were not measured with high confidence due to being poorly represented in the pseudovirus library. The lines above the heatmap indicate different regions of spike. See https://dms-vep.org/229E_spike_1984_DMS/APN_binding.html for interactive plots showing the effects of mutations on hAPN binding.

**Supplementary Figure S6.**
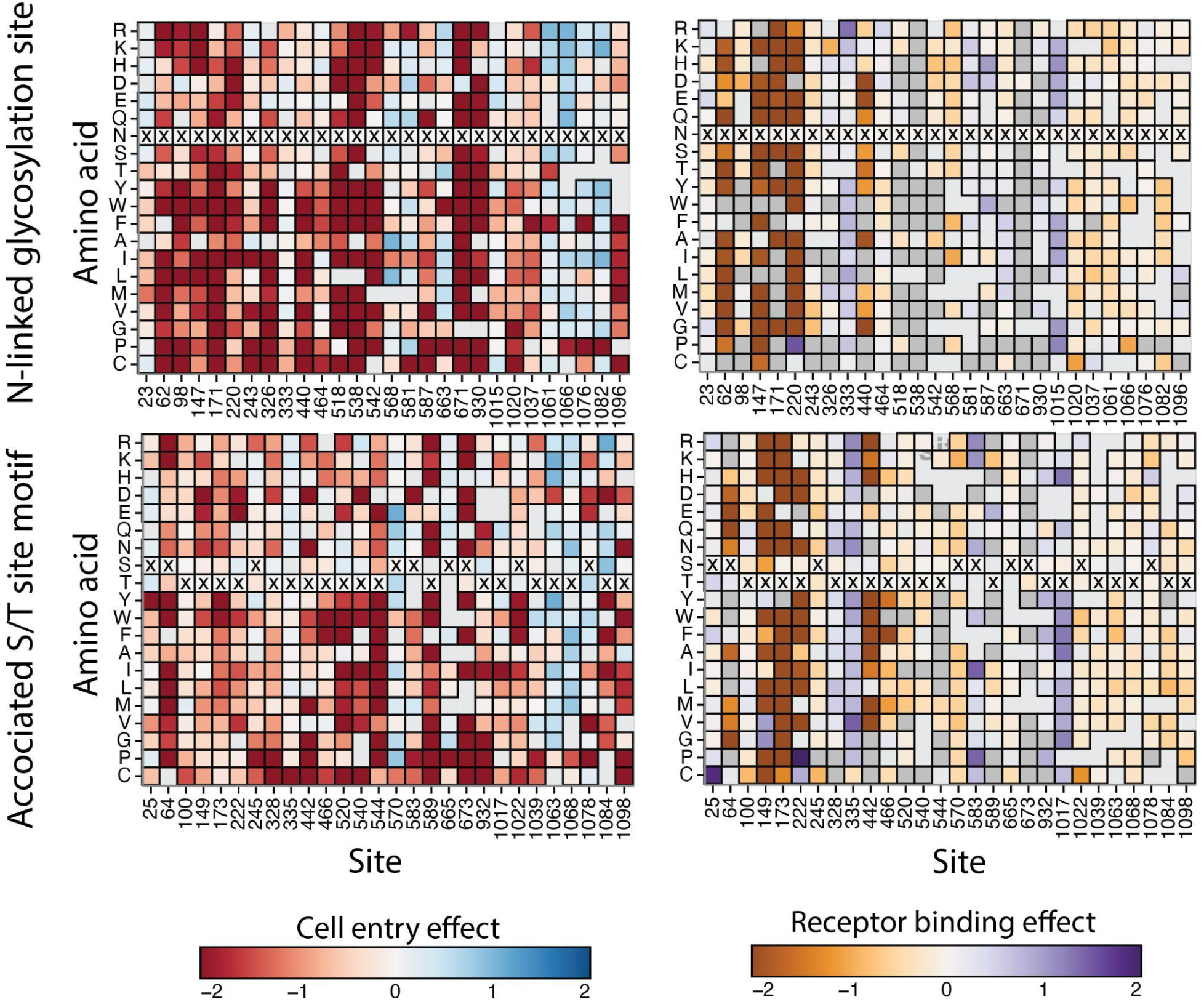
Effects of mutations at N-linked glycosylation motifs on HCoV-229E spike cell entry and hAPN binding. Heatmaps show the effects of amino-acid mutations at N-linked glycosylation motifs (N-X-S/T) and the associated motif positions on spike-mediated cell entry (left) and hAPN binding as measured by pseudovirus neutralization by soluble hAPN (right). Each square represents the effect of an amino-acid mutation, with colors indicating the direction and magnitude of the effect, as in the corresponding full heatmaps for cell entry (Fig. S3) and hAPN binding (Fig. S5). Dark gray squares indicate mutations that strongly impair cell entry (cell entry effect ≤ −2.5) and therefore cannot be reliably measured for their impact on receptor binding; light gray squares indicate mutations that were not measured with high confidence due to poor representation in the pseudovirus library.

**Supplementary Figure S7.**
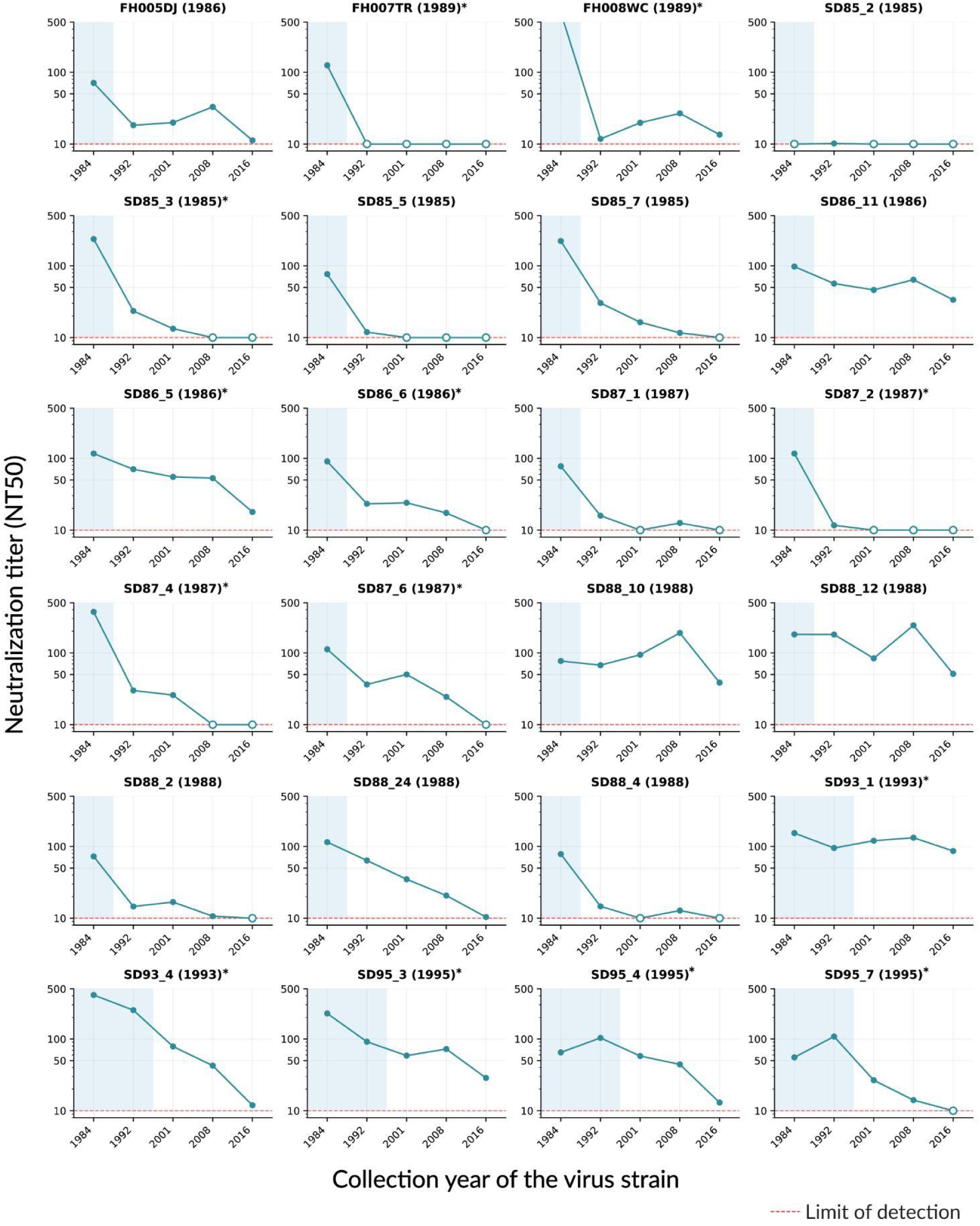
Neutralization of pseudoviruses with HCoV-229E spikes from different years by all tested human sera. This plot is similar to Fig. 5 except it shows all historical human sera tested against the panel of HCoV-229E spikes from different years, whereas Fig. 5 just shows a subset of sera exemplifying either evolution-sensitive or evolution-resistant neutralization patterns.

**Supplementary Figure S8.**
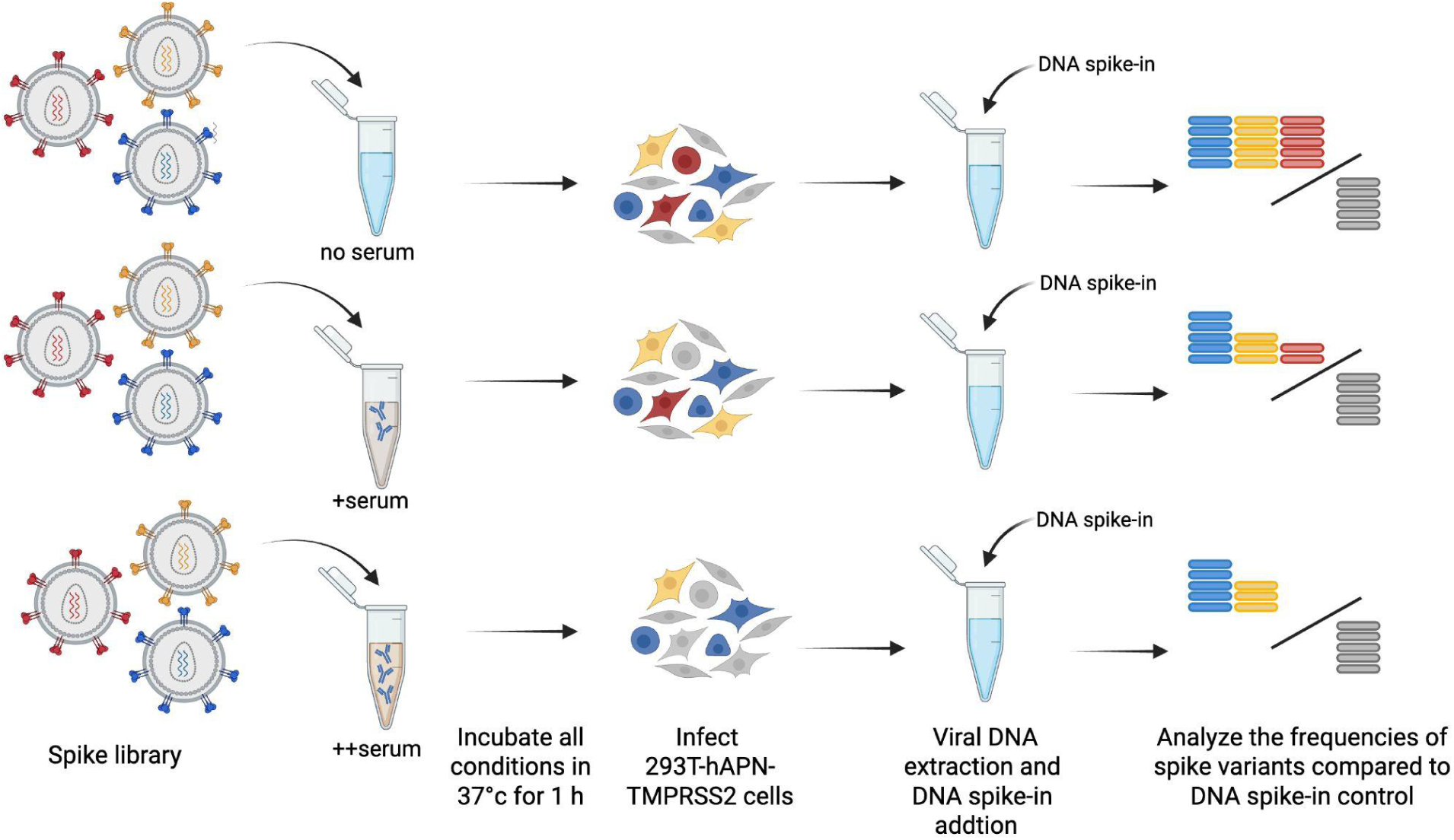
Measurement of effects of spike mutations on serum neutralization escape. The spike-pseudotyped variant library is incubated with increasing concentrations of serum, including a no-serum control. After 1 h incubation at 37°C, 293T-hAPN-TMPRSS2 cells are infected with the pseudovirus mix. At 12 h post-infection, a DNA spike-in is added, and viral DNA is extracted from infected cells. Variant barcode frequencies are quantified by sequencing^29^, and DNA spike-in counts are used to normalize spike-variant read counts across conditions, enabling calculation of the absolute fraction infectivity of each spike variant that is retained at each serum concentration. The fraction infectivities are analyzed to determine the effect of each mutation on serum neutralization.

